# Reconstruction of Contemporary Human Stem Cell Dynamics with Oscillatory Molecular Clocks

**DOI:** 10.1101/2021.03.15.435426

**Authors:** Calum Gabbutt, Ryan O Schenck, Dan Weisenberger, Christopher Kimberley, Alison Berner, Jacob Househam, Eszter Lakatos, Mark Robertson-Tessi, Isabel Martin, Roshani Patel, Sue Clark, Andrew Latchford, Christopher Barnes, Simon J Leedham, Alexander RA Anderson, Trevor A Graham, Darryl Shibata

## Abstract

Molecular clocks record cellular ancestry. However, currently used clocks ‘tick too slowly’ to measure the short-timescale dynamics of cellular renewal in adult tissues. Here we develop ‘rapidly oscillating DNA methylation clocks’ where ongoing (de)methylation causes the clock to ‘tick-tock’ back-and-forth between methylated and unmethylated states like a pendulum. We identify oscillators using standard methylation arrays and develop a mathematical modelling framework to quantitatively measure human adult stem cell dynamics from these data. Small intestinal crypts were inferred to contain slightly more stem cells than colon (6.5 ± 1.0 vs 5.8 ± 1.7 stem cells/crypt) with slower stem cell replacement in small intestine (0.79 ± 0.5 vs 1.1 ± 0.8 replacements/stem cell/year). Germline *APC* mutation increased the number of replacements per crypt (13.0 ± 2.4 replacements/crypt/year vs 6.9 ± 4.6 for healthy colon). In blood, we measure rapid expansion of acute leukaemia and slower growth of chronic disease. Rapidly oscillating molecular clocks are a new methodology to quantitatively measure human somatic cell dynamics.

## Introduction

The fates of individual human cells *in vivo* are difficult to reconstruct. In animal models, the use of transgenic or exogeneous cell labelling enables straightforward clonal lineage tracing (Monné *et al*., 2005; Lopez-Garcia, Allon M. Klein, *et al*., 2010; Snippert *et al*., 2010, 2014; Blanpain and Simons, 2013; Sánchez-Danés *et al*., 2016; Aragona *et al*., 2017; Lan *et al*., 2017; Andersen *et al*., 2019; Han *et al*., 2019), but in humans these methods are precluded. Instead, human studies must utilize somatic genomic alterations, termed ‘molecular clocks’, to trace somatic cell fates. The idea is that ancestry of a population of cells is revealed by the somatic alterations shared amongst the cells: closely related cells are likely to share multiple alterations, whereas distantly related cells will have few alterations in common. In other words, human lineage tracing studies leverage the notion that the clonal history of a cell is recorded in its genome. Various types of somatic genomic alterations have been exploited for lineage tracing, including mitochondrial DNA (mtDNA) mutations (Taylor *et al*., 2003; Greaves *et al*., 2006; Fellous *et al*., 2009; Gutierrez-Gonzalez *et al*., 2009; Gaisa, Graham, McDonald, Cañadillas-Lopez, *et al*., 2011; Gaisa, Graham, McDonald, Poulsom, *et al*., 2011; Humphries *et al*., 2013; Baker *et al*., 2014, 2019; Lavery *et al*., 2014; Moad *et al*., 2017; Stamp *et al*., 2018; Cereser *et al*., 2018; Ludwig *et al*., 2019), DNA methylation at neutral loci (Yatabe, Tavaré and Shibata, 2001; Kim and Shibata, 2004; Kim *et al*., 2005; Kim, Tavaré and Shibata, 2006; Chu *et al*., 2007; Nicolas *et al*., 2007; Siegmund *et al*., 2009; Graham *et al*., 2011), allelic loss at heterozygous loci (Campbell *et al*., 1994; Novelli *et al*., 2003) and the detection of single nucleotide variants (SNVs) via genome sequencing (Leedham *et al*., 2008, 2009; Thirlwell *et al*., 2010; Galandiuk *et al*., 2012; Pipinikas *et al*., 2014; Martincorena *et al*., 2015; Blokzijl *et al*., 2016; Williams *et al*., 2016, 2018; Simons, 2016; Lee-Six *et al*., 2018; Caroline J. Watson *et al*., 2020; Moore *et al*., 2020).

These molecular clocks use ‘unidirectional’ measurements that essentially count the accumulation of changes since birth to infer the relatedness between lineages. The resolution at which a molecular clock can track clonal ancestry is a function of the rate at which genomic alterations accrue. A low rate of alteration accrual (situations where the molecular clock ‘ticks’ slowly) reveal clonal dynamics occurring over long timescales. For example, genome sequencing studies of normal skin (Martincorena *et al*., 2015), blood (Lee-Six *et al*., 2018), intestinal crypts (Blokzijl *et al*., 2016), and endometrial glands (Moore *et al*., 2020) identified multiple subclones in each tissue, but in most cases reconstructed lineages diverged many years in the past and recent cell turnover was not evident in the data. In comparison, a fast rate of alteration accrual (situations where the clock ticks rapidly) has the potential to reveal rapid and/or recent clonal dynamics, but in practice the clock becomes compromised by ‘saturation’ wherein the same pattern of alterations convergently evolve in distinct clonal populations (Kuipers *et al*., 2017), and effectively recording stops in childhood.

Here we introduce the concept of a new class of *rapidly oscillating molecular clocks* where, like a pendulum, genomic alterations can reversibly change their states (the clock ‘tick-tocks’ back and forth). We propose that certain CpG sites rapidly and stochastically oscillate their DNA methylation between 0% (homozygously unmethylated CpG), 50% (heterozygous methylation), and 100% (homozygous methylation) in individual diploid cells (Fig. 1A). When this oscillation occurs at a timescale similar to the cell division rate, we show that measurements of these oscillators can be used to infer contemporary cell population dynamics. The analysis of oscillator clocks is more complicated than unidirectional (“hourglass”) clocks. However, the rapid ‘tick-tock’ (methylation-demethylation) alterations facilitate the study of contemporary events that occur later in life, or indeed which recur throughout life.

**Figure 1:**
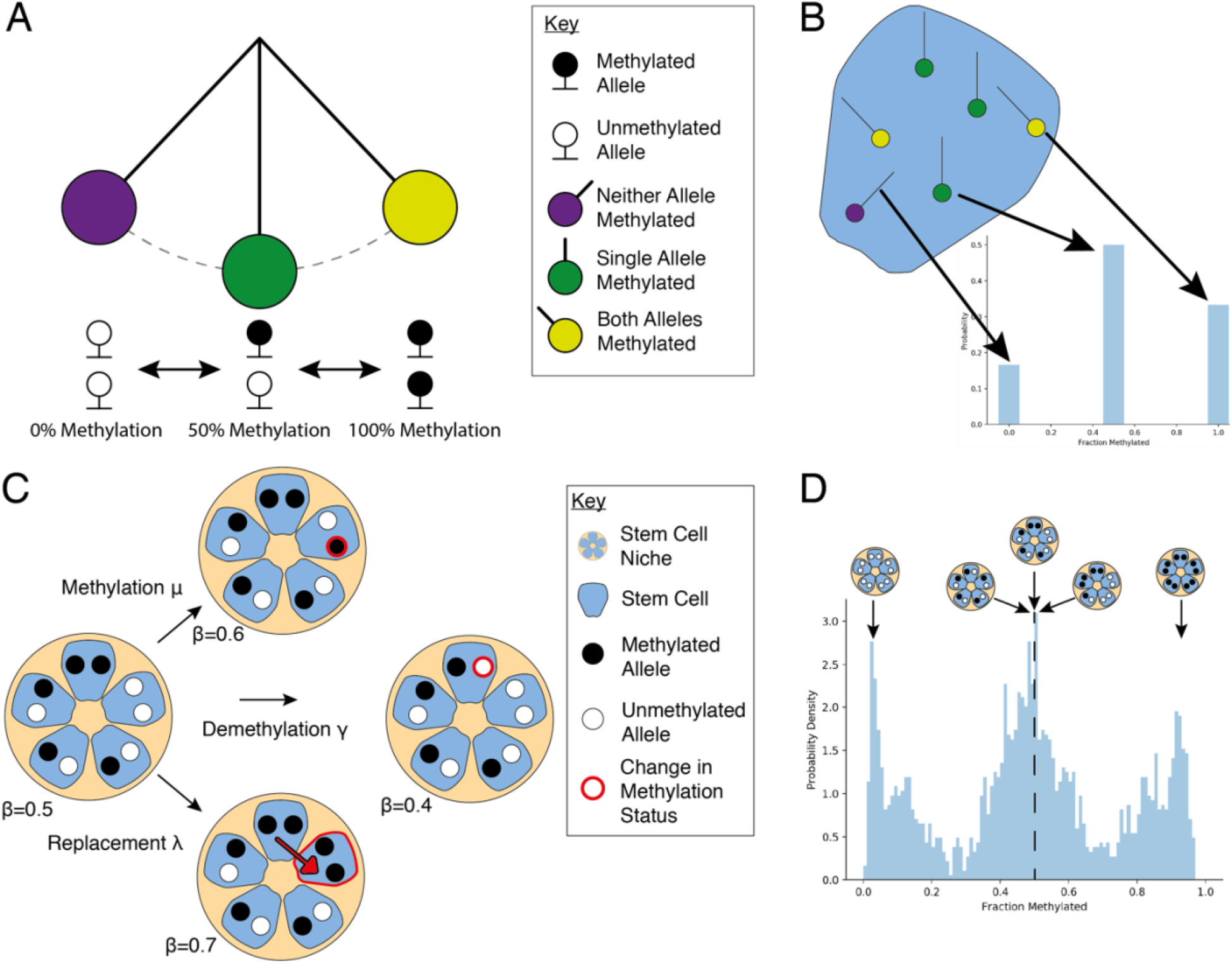
The Tick-Tock Clock – Oscillatory Methylation as a Lineage Tracing Marker. **A:** Illustration of the three possible methylation states at a specific CpG locus within a particular cell. A cell can either be homozygously (de)methylated, or heterozygously methylated at that CpG locus. It is the spontaneous transitions between these states that allow methylation to act as a lineage tracing marker. **B:** Illustration of the link between the methylation status of a given CpG locus within a particular cell and the beta value (the fraction of methylated DNA at that locus) associated with that cell. **C**: Graphical representation of how the methylation status in a small population of 5 stem cells at a particular CpG locus can change over time due to (i) methylation, (ii) demethylation, or (iii) cell replacement. **D:** Methylation (beta) distributions from an individual crypt, the peaks near 0% and 100% correspond to a clonal methylated or unmethylated CpG loci respectively, whereas the peak at 50% corresponds both to clonal heterozygous CpG loci and subclonal populations caught mid-sweep.

Here we show how ‘tick-tock clock’ CpG oscillator methylation can be conveniently measured with commercial microarrays (Illumina EPIC arrays) that provide the methylation value at thousands of candidate CpG clocks. We develop a mathematical inference methodology to extract ancestral information encoded within oscillating sites. We validate our methodology using a simplified spatial model of a crypt cell evolution driven by different stem cell numbers then apply our tick-tock clock methodology to measure stem cell dynamics in individual human intestinal crypt and endometrial gland populations. The oscillators are further applied to whole blood to detect and distinguish between acute and chronic leukaemias.

## Results

Here, we present evidence that a set of CpG loci rapidly oscillate their methylation status and can be used to perform lineage tracing of recently and/or rapidly occurring clonal expansions in human tissues. We apply the new method to quantify the dynamics of clonal expansions in human colon, small intestine, endometrium and blood.

### Identification of Oscillatory CpG Loci

We isolated DNA from individual single colon or small intestinal (SI) crypts (31 colon samples originating from 8 patients, and 28 SI samples originating from 7 patients) and measured DNA methylation in each crypt using Illumina EPIC arrays (methods). Samples from each tissue were treated separately to account for tissue-specific differences in DNA (de)methylation processes.

To select oscillating CpG sites, we first removed ∼400,000 CpG loci from the ∼850,000 on the EPIC array that were likely to be actively regulated or associated with probes that co-hybridize highly homologous sequences (methods). CpG sites with oscillating methylation were then detected by comparing between-patient versus within-patient heterogeneity in methylation value. At rapidly oscillating sites, we expect the average methylation in non-clonal ‘bulk’ samples to be 0.5 (since methylation at the site is uncorrelated between the multiple lineages that make up the bulk sample), whereas in individual clonal samples the methylation value can take any value between 0 and 1. Thus, to select for oscillating CpGs, we selected CpG sites that had the highest 5% of variance in beta value between individual samples, and then filtered these for sites with mean methylation across all samples and patients of ∼0.5 (mean beta value between 0.4 and 0.6) (Fig. 2A). This process identified 7073 putative oscillatory CpGs within the colon sample cohort and 8828 CpGs within the small intestine cohort, of which 1794 CpGs were shared between tissue types (Fig. S1A). There was a good correlation (R^2=0.62) in the heterogeneity scores between colon and small intestine samples (Fig. S1B), and CpG loci that were exclusive to the colon had a substantially higher average variability score in the small intestine compared to all CpG loci (Fig. S1C), suggesting that the relatively large number of non-overlapping loci was due to the arbitrary strictness of our 5% threshold. Further analysis was performed upon these shared 1794 oscillatory CpG loci (see supplementary table 1) with the goal of aiding the generalizability of our approach. The 1794 oscillators had ∼50% average methylation among the 31 crypts, but exhibited a characteristic “W-distribution” of methylation values within the recently clonally-derived population of cells that make up an individual crypt (Fig. 1D).

**Figure 2:**
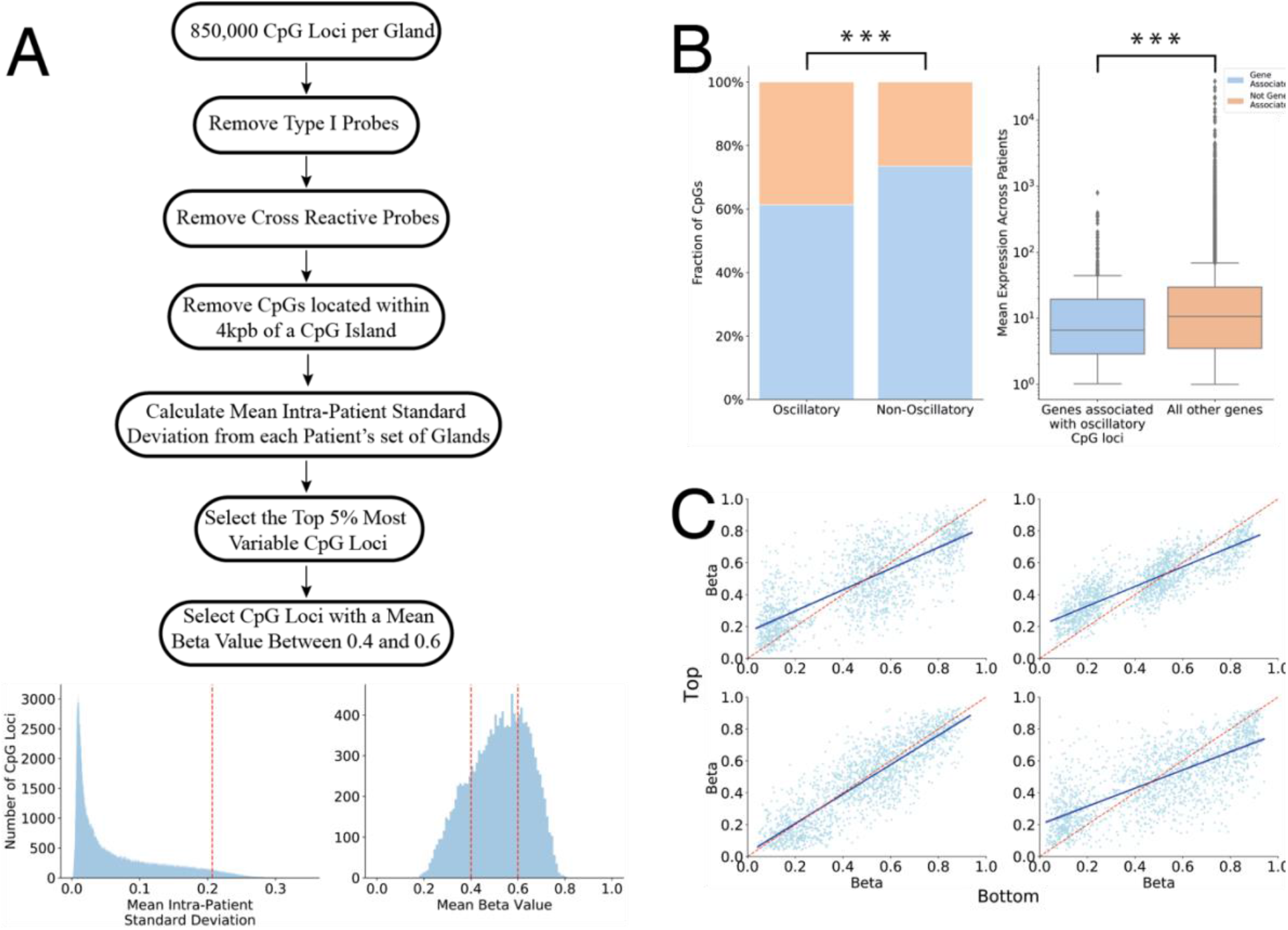
Identification of Selectively Neutral Oscillatory CpG Loci. **A:** Workflow used to identify oscillatory CpG loci that exhibit high intra-patient heterogeneity. Input data was the ∼850,000 CpG loci assayed by an Illumina EPIC array. We removed Type I probes and probes which cross-hybridize highly homologous DNA regions. For each CpG locus, we calculated the standard deviation for each set of ∼4 crypts per patient, and then calculated the mean standard deviation across the patient cohort as a metric for the intra-patient heterogeneity. We selected the top 5% most highly variable CpG loci, and then removed CpG loci which have a mean beta value (across the entire patient cohort) less than 0.4 or greater than 0.6. **B:** (left) Oscillatory CpGs are enriched for CpG loci not associated with any genes. (right) The set of genes associated with the oscillatory CpGs exhibit a lower average RNA expression in normal colon than those genes associated with non-oscillatory CpG loci. **C:** Beta values of oscillatory CpG loci are correlated between the bottom and top halves of a crypt.

To demonstrate technical accuracy in methylation measurement from the small amounts of DNA in single SI crypts (∼400 cells), colon crypts (∼2,000 cells) or endometrial glands (∼5,000 cells), we identified similar oscillators on the X-chromosome and compared methylation between male and female individuals. In males, there is only a single copy of the X-chromosome, hence only two modal peaks near 0 and 100% methylation should be present in clonal populations, as opposed to the trimodal distribution observed on autosomes. Consistent with the ability to measure oscillator methylation in small tissue samples, the X-chromosome oscillators exhibited W-shaped distributions in female SI crypts and “U-shaped” distributions in male SI crypts (Fig. S1D).

### Oscillatory CpG Loci Are Enriched in Minimally Expressed Genes

For CpG loci to act as a molecular clock, the loci must not be subject to evolutionary selection or cell-specific regulation. We compared the proportion of the 1794 oscillatory CpG sites that were associated with a specific gene to the 428511 CpG sites that were not identified as oscillatory (methods). Oscillatory CpG loci were strongly enriched for non-genic CpG sites (Fig. 2B 1.8 OR, chi-squared test, p<0.001). We tested RNA expression using 40 normal colon samples from TCGA (Muzny *et al*., 2012) and found that the mean expression of genes associated with oscillatory CpG loci was lower than of genes associated with the non-oscillatory CpG loci (Fig. 2B, -0.24 Cohen’s d calculated for log-transformed expression, Welch’s unequal variance t-test, p<0.001). Together, these analyses indicated that methylation at the oscillatory CpG sites was unlikely to be under strict regulation or evolutionary selection in the colon.

### Methylation status of oscillatory CpG loci are preserved along the length of the colon crypt

Previous research (Kaaij *et al*., 2013) has found that the methylation profile of the whole crypt was, on the whole, representative of that of the stem cell population at the base of the crypt. To ensure that this was the case specifically for the oscillatory CpG loci identified above, we split 7 crypts into their respective tops and bottoms, and ran Illumina EPIC arrays upon both halves using the same protocol described previously. Due to the low input DNA amounts, 3 of the samples failed the QC step. The remaining 4 crypts exhibited a good correlation between the beta values of the oscillatory CpG loci in the tops and bottoms of the crypts (Fig. 2C, R^2^ > 0.6, p < 0.001 in all cases).

### Mathematical Model of Stem Cell Evolution and the Beta Distribution of Oscillatory CpG Loci

We hypothesized that the precise shape of the methylation beta-value distribution for oscillatory CpGs was determined by the underlying dynamics of cellular evolution. To test this hypothesis in the context of intestinal crypts, we developed a mathematical model and associated Bayesian inference framework to relate the competitive dynamics of stem cells within their crypt to the measured distribution of oscillatory CpG methylation.

The mathematical model consisted of a hidden Markov chain that simulated the time-dependent probability distribution of the number of methylated and unmethylated copies of a single CpG locus within a stem cell niche of fixed size *S*. We described 3 possible processes that changed the methylation status at a given CpG locus: (1) spontaneous methylation (at constant rate *μ* per allele per stem cell per year), (2) spontaneously demethylation (constant rate γ per allele per stem cell per year), and (3) one stem cell replacing another stem cell (constant rate λ per stem cell per year) (Fig 1C). We further assumed that the stem cells could be treated as a well-mixed population, such that each stem cell could replace any other stem cell within the niche with equal probability. The probability distribution of the methylation value at a single CpG site could be fully characterized with just two state variables: *k* the number of stem-cells in the crypt with just one allele methylated, and *m* the number of stem cells with both alleles methylated. By considering the possible (*k, m*) → (*k*^′^, *m*^′^) transitions, we derived a system of ordinary differential equations describing how the probability (p(*k, m*|*λ, μ, γ*; *t*)) of the system being in state (*k, m*) changes over time (see the methods section for an overview and the appendix for a full derivation). For a pool of *S* stem cells, there are 2*S* + 1 discrete states the niche methylation level could take, with a beta value of 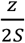 (for *z ∈ [*0, 2*S]*). To link the probability that a particular CpG locus has a population methylation status *z* to the output of our stem cell dynamics model, we marginalised over the various combinations of *k* and *m* that correspond to a particular *z*-value, as described in the methods section.

We developed a Bayesian inference framework (Methods), that allowed for simultaneous inference of the number of stem cells (*S*), the replacement rate per stem cells (*λ*), and the methylation (*μ*) and demethylation (γ) rates per stem cell per allele per year for an individual patient-derived gland. This Bayesian pipeline accounted for the error profile of the methylation array technology, such as the observed ‘offset’ from 0% and 100% methylation owing to background noise, and the uncertainty in the methylation (beta value) measurement due to technical noise in the assay and the noise generated by sampling a limited number of alleles for analysis. Thus, we could fit our model of stem cell dynamics to the data from individual crypts, allowing us to probe tissue-specific stem cell dynamics whilst accounting for intra- and inter-patient heterogeneity.

### Evolutionary Dynamics Are Inferred with High Accuracy *In Silico*

To verify that our Bayesian inference framework was able to accurately infer the stem cell dynamics of a crypt, we generated three “synthetic” crypts each containing 5 stem cells, a mean replacement rate of 1 per stem cell per year and a *de novo* (de-)methylation event rate of 0.0005, 0.05 and 0.5 per allele per stem cell per year (Fig. 3A) and used our inference framework to attempt to recover the (known) underlying parameter values from the simulated methylation distributions.

**Figure 3:**
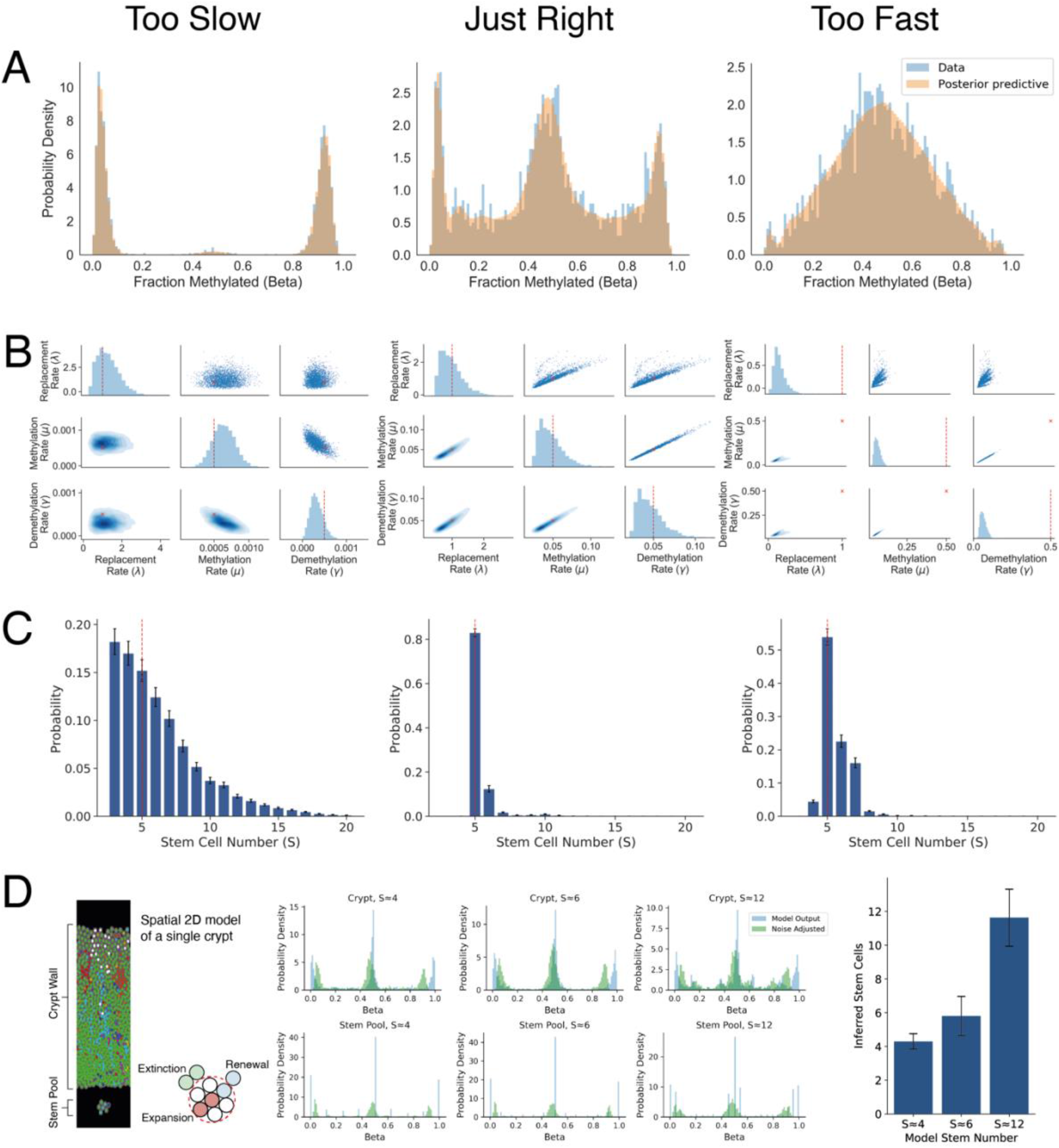
W-Shaped Beta Distributions Are Indicative of Informative Clonal Information. *In silico* evaluation of the accuracy of Bayesian inference on stem cell number (*S*), replacement rate (*λ*) and (de)methylation rates (*μ, γ*) as a function of input (de)methylation rates. Three regimes were evaluated: *μ* = *γ* = 0.0005 methylation events per allele per stem cell per year, termed “too slow”, very high methylation rates *μ* = *γ* = 0.5 per allele per stem cell per year, termed “too fast”, and intermediate methylation rates (*μ* = *γ* = 0.05 per allele per stem cell per year, termed “just right”. **A**: Simulated oscillatory CpG methylation distributions from individual crypts at each of three input (de)methylation rates. The characteristic W-distribution is only evident for the “just-right” (de)methylation rate. **B:** Posterior distributions of inferred replacement and (de)methylation rates for each input (de)methylation rate. **C:** Posterior distributions of inferred stem cell number. In panels B and C, red dashed lines indicate the true (inputted) value of the parameter. The simulated datasets each contained *S =* 5 stem cells, had a replacement rate of *λ* = 1.0 per stem cell per year, and the noise due to sampling was simulated with offsets due to background noise Δ= *ε* = 0.05 and peak specific noise with sample size *k*_*z*_ = 100. **D:** Independent validation of the inference method on a spatial representation of the single crypt with varying stem cells. Beta distributions are noise adjusted (methods) for the inferences on the stem pool only. Mean inferred stem cells are shown for ten replicate simulations, error bars denote standard deviations.

At low (de)methylation rates (where the tick-tock clock oscillated “too slowly”), the methylation distribution was essentially concentrated near 0% and 100% methylated, with a small minority of CpG loci in the intermediate 50% methylation state, mainly due to clonal heterogeneous methylation. Conversely, a high (de)methylation rates (where the tick-tock clock oscillated “too fast”) the methylation distribution approached a binomial-like distribution centered at 50%. Unlike the “too slow” crypt, the peak at 50% is largely due to sub-clonal mutations caught mid-drift, rather than a single fixed mutation. At intermediate (de)methylation rates (where the tick-tock clock oscillated at a “just right” rate) crypt methylation distributions showed the same characteristic W-shape that we observed in the real patient crypt data. Major peaks were evident near 0%, 50% and 100%, and additional minor peaks ∼10%-40% and ∼60-90%, which are due to methylation events that had not fixed (sub-clonal (de)methylation events).

Bayesian inference could not satisfactorily determine the posterior for the number of stem cells for the “too slow” crypt, as there were too few CpG sites with intermediate values that held information on stem cell number. In contrast, the inference framework accurately recovered the number of stem cells for the “too fast” crypt, as there was an abundance of sub-clonal methylation events, but the replacement rate could not be inferred accurately. This is because the clonal information that is propagated by stem cell replacement (increase/decrease in beta values from the expanding clone) is almost immediately lost due to the high (de)methylation rate.

When the simulated methylation rate was “just right” the model was accurately able to recover all known parameter values with good confidence (Fig. 3B & 3C). We note that this *in silico* analysis shows that we are able to confidently confirm that the (de)methylation rate for a given set of CpG loci is within the “just right” range by the presence of the characteristic W-shape. Note that the range of the methylation error rates that give rise to the W-shape and which are suitable for timing using our analysis is relatively broad, covering over 2 orders of magnitude.

To further validate our Bayesian inference framework, we implemented a simplified agent based spatial model of crypt cell evolution (methods) where each cell (agent) incorporates molecular level CpG tracking with (de)methylation errors possible upon each cell division. We used this *in silico* crypt model to generate tick-tock CpG patterns from a range of stem cell pool sizes. Then applying our inference framework on the resulting CpG patterns we were able to accurately recover the stem cell numbers (Fig. 3D), for each of the three different pool sizes (3.76±0.73, 6.42±0.98, and 12.39±1.16 stem cells; mean ± standard deviation).

### Measurement of Stem Cell Dynamics in Human Intestine

We applied our inference framework to the distribution of methylation values observed across oscillatory CpG loci from human colon and small intestinal crypts. Fits were performed on each crypt and patient individually, generating crypt and patient-specific posterior estimates of effective stem cell number and replacement rate (Fig 4A).

**Figure 4:**
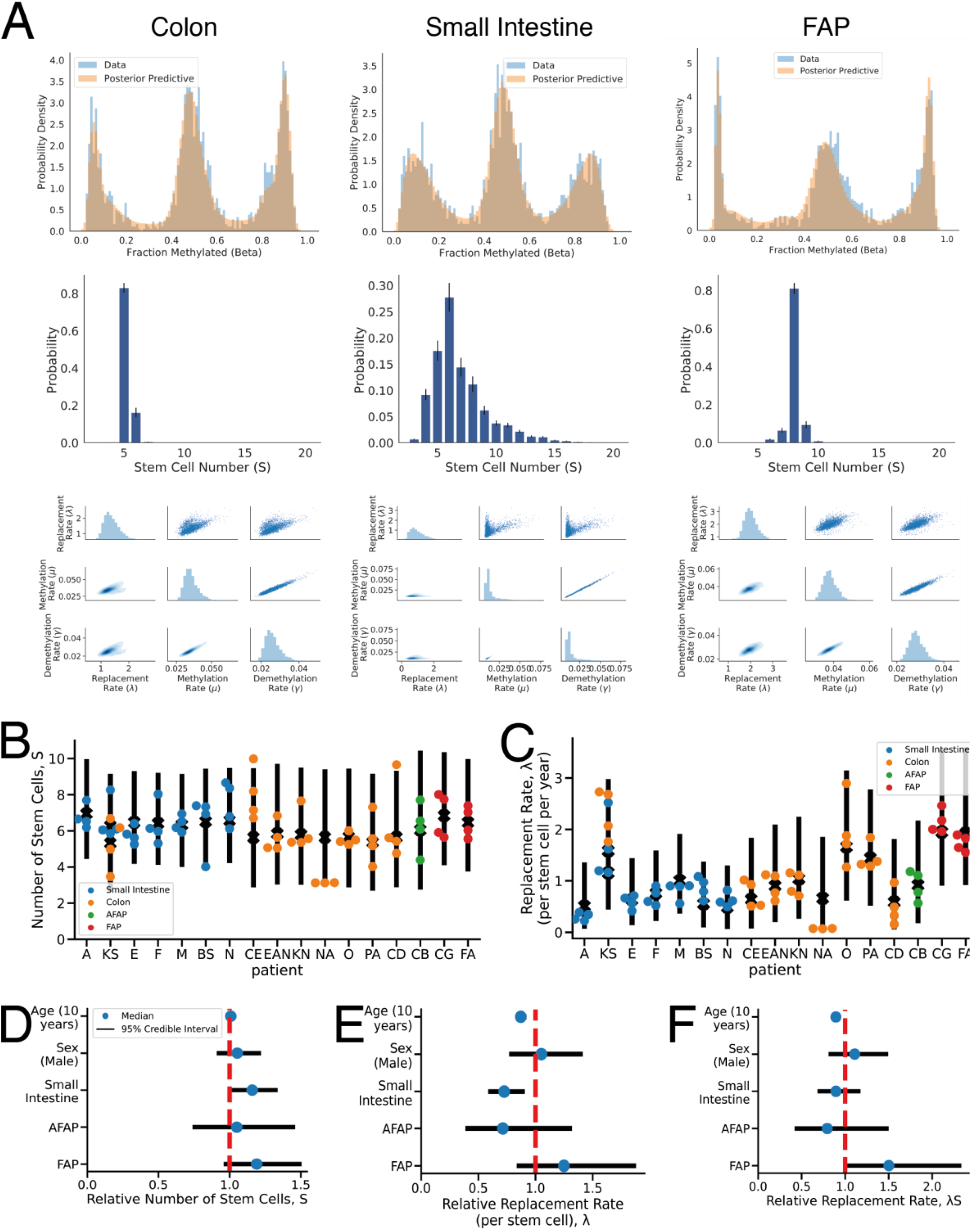
Tissue Specific Differences in the Stem Cell Dynamics. **A:** Examples of the posterior predictive distributions, the discrete stem cell number posterior, and the posterior for the replacement rate, methylation rate and demethylation rate in crypts derived from normal colon, small intestine and the colon of patients with FAP (left to right panels). **B**: individual crypt and posterior mean per patient for the stem cell number and **C:** replacement rate per stem cell, with the 95% credible range of the generalized linear model (GLM) expectation, accounting for age, sex, tissue, disease state and intra- and inter-patient heterogeneity. **D-F**: posterior distributions for the effect of patient age (per decade), sex (with female encoded as reference), tissue type and disease state on the relative number of stem cells, replacement rate per stem cell and total number of replacements when compared to normal colon. A Bayesian parameter estimation hypothesis testing approach was taken, such that a difference was called significant if the 95% credible region did not overlap 1.

The mean number of stem cells was similar across tissues, with 5.8 ± 1.7 stem cells in normal colon samples, and 6.5 ± 1.0 stem cells within small intestinal glands (mean ± 1 standard deviation, Fig. 4B). The replacement rate in normal colon was 1.1 ± 0.8 replacements/stem cell/year, reduced to 0.79 ± 0.5 replacements/stem cell/year in small intestine (Fig. 4C).

We used a hierarchical Bayesian generalized linear model (GLM) to account for patient structure in our data and compared stem cell numbers and replacement rates between tissues. We found that glands from the small intestine had a greater number of stem cells (Fig. 4D, p<0.05; GLM), but a lower replacement rate per stem cell compared to normal colon (Fig. 4E), such that the total number of replacements per crypt was not significantly different between colon and small intestine (Fig. 4F, p<0.05; GLM).

Patients with familial adenomatous polyposis (FAP) carry a heterozygous germline mutation in the *APC* gene and are increased risk of developing colorectal cancer (Groden *et al*., 1991; Kinzler *et al*., 1991; Nishisho *et al*., 1991). APC is a key regulator of wnt-signalling, and consequently pathogenic *APC* mutations cause alterations to wnt-signalling (Korinek *et al*., 1997; Sansom *et al*., 2004; Zhan, Rindtorff and Boutros, 2017). Wnt-signalling is essential for maintenance of intestinal stem cells (Korinek *et al*., 1998; Pinto *et al*., 2003; Pinto and Clevers, 2005). Consequently, we hypothesized that FAP patients would have altered stem cell dynamics. Inference on oscillating CpG sites showed that stem cell numbers were similar in FAP crypts to normal colon (6.7 ± 0.3 stem cells per crypt), but the stem cell replacement rate was almost doubled at 1.9 ± 0.3 replacements/stem cell/year (Fig. 4A-C), resulting in a significantly higher total number of replacements per crypt per year in FAP (Fig. 4F).

### Stem Cell Dynamics in Human Endometrial Glands

We analyzed tick-tock rapidly oscillating CpG methylation in 32 endometrial glands derived from 8 patients using the same methodology as for intestinal crypts (Fig. 5). We derived a set of 7721 oscillatory CpG sites, of which 807 were shared with the set of oscillatory loci identified in the colon. The resulting methylation beta distributions exhibited the same characteristic W-shape as in the intestine (Fig. 5A).

**Figure 5:**
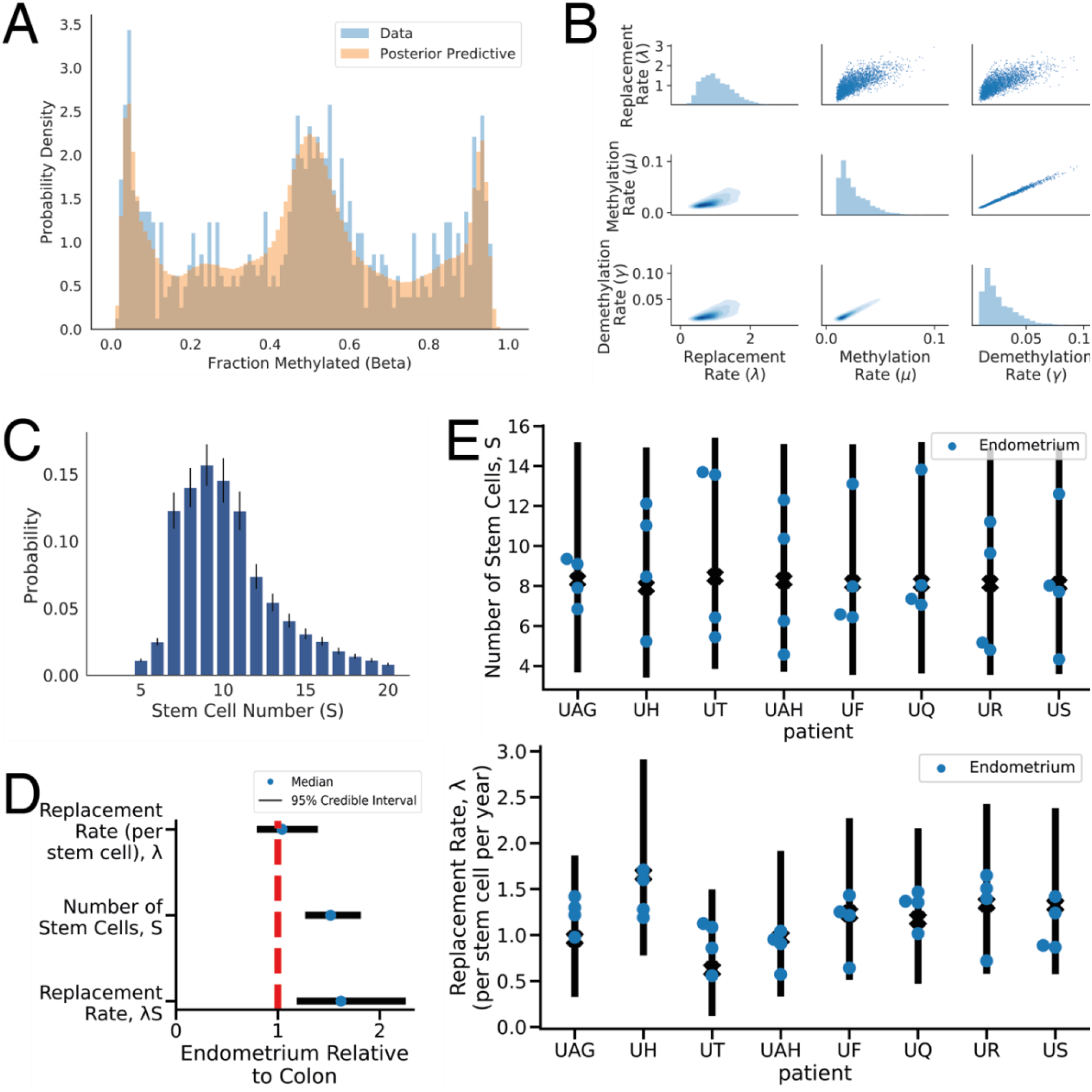
The Tick-Tock Method Is Generalizable to Other Glandular Tissue. **A:** Measured methylation beta values (blue bars) and posterior predictive distribution (salmon overlay) for an example endometrial gland. The methylation patterns exhibit a similar W-shape to that observed in intestinal crypts. **B**: Posterior distributions for the replacement rate per stem cell and (de)methylation rates for the gland shown in (A). **C**: Posterior distribution for stem cell number for the gland shown in (A). **D:** Inferred relative replacement rate per stem cell, number of stem cells and total replacement rate, in endometrium versus colon, indicating that there are significantly more stem cells per gland in endometrium than colon. Bars showed 95% credible intervals derived from a GLM. **E**: Inferred number of stem cells and replacement rate per stem cell for each individual gland from each patient (dots) with the 95% credible range of the GLM expectation (bars).

We then applied our Bayesian inference pipeline to each endometrial gland to infer the effective stem cell dynamics (Kim, Tavaré and Shibata, 2005). The inferred stem cell replacement rate was broadly similar compared to colon at 1.2 ± 0.3 (mean ± 1 standard deviation) replacements/stem cell/year (Fig. 5B), whereas the number of stem cells per gland was significantly higher in endometrium compared to colon (p<0.05; GLM), with each endometrial gland containing 8.6 ± 2.9 stem cells (Fig. 5C). Intriguingly, the endometrium exhibited a significantly greater degree of intra-patient variability with regards to the number of stem cells (p<0.05; GLM), perhaps due to the dynamic nature of the endometrium through menstrual cycles and age-related changes. We acknowledge that the stem cell structure of endometrial glands is likely more complex than that of colon crypts (Tempest *et al*., 2020), limiting the degree to which our simple model reflects the underlying biology. Nevertheless, the fact that we still observe large clonal peaks near 0% and 100% methylation suggests that monoclonal conversion does still occur, and our model is still applicable as a simplified caricature of the complicated dynamics present in endometrial glands.

### Tick-Tock Clock in Human Blood

The CpG oscillator behavior seen in intestinal crypts and endometrial glands are likely to be present across tissues. Therefore, we next searched for similar oscillators in whole human blood, which has abundant public methylation array data for normal and disease states. Unlike intestinal crypts which recurrently drift to clonality, blood is a large, well-mixed tissue with diverse cell types and is normally polyclonal because it is produced by thousands of bone marrow stem cells (Lee-Six *et al*., 2018; Caroline J Watson *et al*., 2020). However, blood becomes oligoclonal or clonal in disease states such as leukaemias. As in the intestines, CpG sites that rapidly oscillate through 0, 50 and 100% methylation will have average methylation around 50% in normal polyclonal blood samples.

We identified suitable tick-tock loci by averaging normal whole blood DNA methylation at ∼450,000 autosomal CpG loci from a commonly used aging database of 656 healthy individuals (Hannum *et al*., 2013). We selected all loci (N = 27,634) with average values between 40 and 60% methylation in these 656 specimens. CpG oscillators appear tissue specific because only ∼5% of the intestinal tick-tock loci were in the blood set. Tick-tock methylation for each individual sample revealed tight distributions around 50% methylation, which can be described by its variance (Fig 6A). Consistent with oscillators, average tick-tock methylation remained ∼50% with aging (S1). Serial samples ten years apart (Tan *et al*., 2016) revealed tick-tock variance to be relatively stable for an individual, with a slight significant trend for increases with age (Fig 6B), which was also observed throughout aging (Fig 6A).

**Figure 6:**
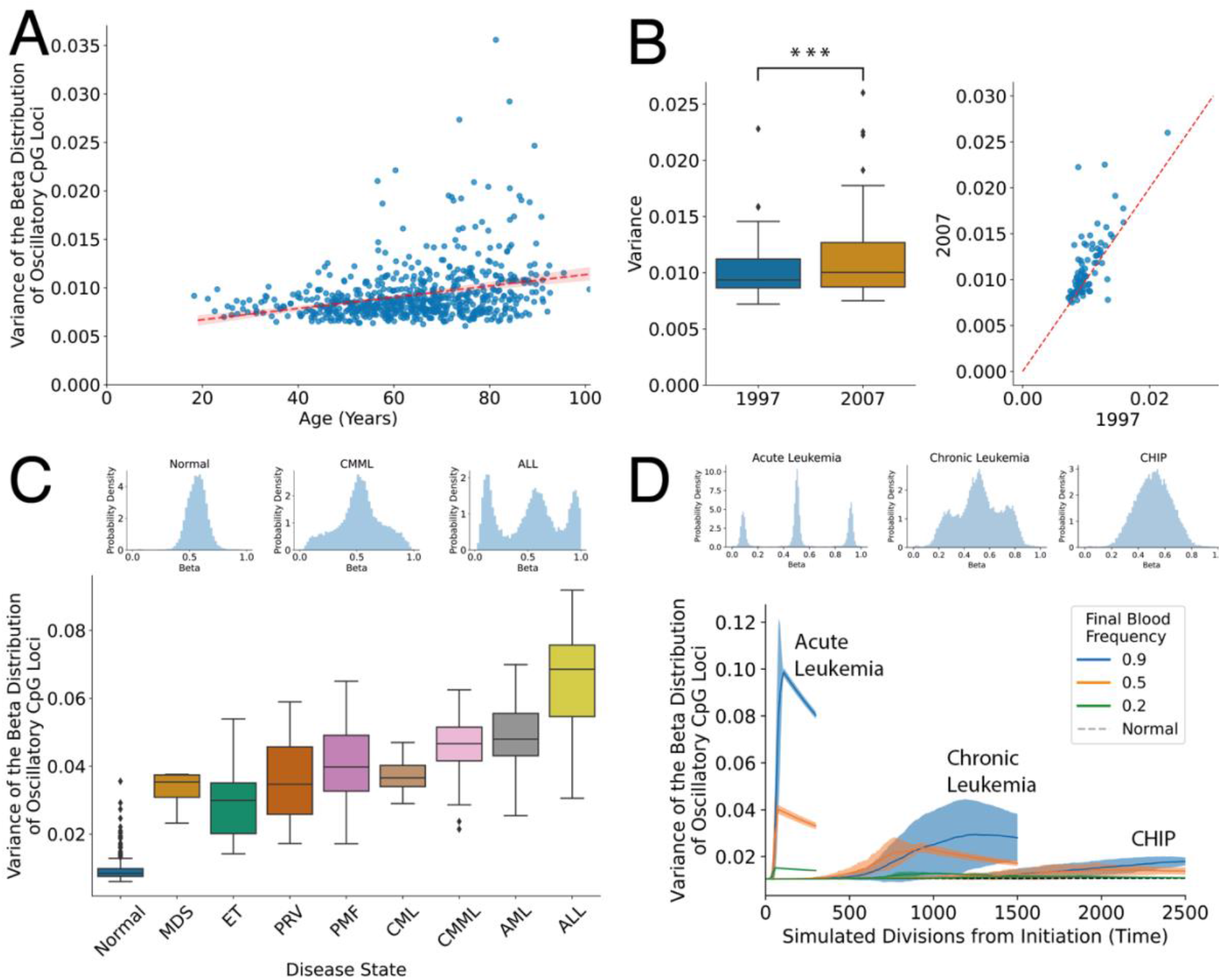
Tick-Tock Dynamics Can Further be Observed in Chronic and Acute Leukemia. **A:** The variance of the oscillatory CpG beta distribution experiences a gradual increase with age in normal patients. **B:** The variance of paired blood samples taken 10 years apart (1997 and 2007) also exhibits a small but marked increase (0.37 Cohen’s d, p<0.001 paired t test). **C:** The variance of the oscillatory CpG beta distribution is a proxy for the rapidity of the clonal expansion within the blood. In normal samples the large stem cell population size leads to the beta distribution being concentrated near 50% (as one would expect for uncorrelated oscillators). However, as a clonal cancerous population expands, clonal peaks begin to separate from the 50% peak. In the case of ALL, the large, well-separated peaks near 0% and 100% are indicative of a single clonal population making up the majority of the remaining stem cells following rapid growth. **D:** Simulations confirm that a simple model of hematopoiesis can recapitulate the observed beta distribution overserved in patient data.

Clonal hematopoiesis in the blood is an early step in the evolution of neoplasia and will increase tick-tock variances. For rapid clonal expansions (i.e. acute leukaemias), W-shaped blood distributions similar to those observed in the crypts are expected. Consistent with these expectations, whole blood samples from different types of major hematopoietic neoplasm had higher than normal tick-tock variances (Fig 6C). Acute lymphoblastic leukaemias (ALL) and acute myeloid leukaemias (AML) had the highest tick-tock variances and characteristic W-shaped distributions. More indolent chronic myeloproliferative or myelodysplastic whole blood specimens showed more modest tick-tock variance increases and generally lacked the “W” shape of the acute leukaemias, crypts and glands.

### Hematopoiesis Simulations

We simulated hematopoiesis to better understand how oscillators detect clonality in whole blood (Fig 6 and Supplemental). Tick-tock methylation oscillates between 0, 50 and 100% in single cells and the simulations indicate polyclonal whole blood variance is low and stable through time because human hematopoiesis is maintained by large numbers of stem cells. Clonal expansion by a single cell synchronizes oscillators and results in higher whole blood variances that depend on growth rates. As in the crypts, there is a balance between clonal expansion rates, which increase population variances, and the rates at which oscillators drift back to 50% average methylation, which decreases variance. A rapid expansion (less than 2 years) to high blood levels as in acute leukemias produces high variances and W-shaped distributions. The “W” methylation pattern resembles the methylation at 0, 50 and 100% methylation of the initiating cell. Expansions that grow more slowly have variances greater than normal blood but lack the W-shape as clonal oscillators become increasing desynchronized with time. These more indolent expansions are more consistent with the experimental data for chronic myeloproliferative neoplasms, which may be asymptomatic and persist for years. Clones that grow even slower and arise later, as may occur with CHIP (Jaiswal and Ebert, 2019), leads to slightly higher variances, as seen with aging in the normal whole blood cohort. Hence, a simple model with 27,634 oscillating CpG sites and different rates of clonal expansion is broadly consistent with the experiment data from hundreds of clinical samples.

## Discussion

Here we discover and demonstrate how to model a novel class of rapidly oscillating CpG methylation molecular clocks that can reconstruct contemporary human cell population dynamics that start or recur at different times during life, using standard Illumina EPIC methylation arrays applied to bulk tissue samples. Large numbers of CpG sites reversibly oscillate their methylation like a pendulum between 0, 50 and 100% (representing homozygous and heterozygous (de)methylation). In polyclonal populations, these oscillations are unsynchronized between individual cells, and average oscillator CpG methylation is around 50%. However, oscillator methylation that occurs at similar rate to the rate of cell division and replacement within a clonal population, leads to a characteristic W-shaped distribution with modal peaks at 0, 50 and 100% methylation for each CpG site upon bulk measurement of the clone that resemble the ‘tick-tock’ state of the most recent common ancestor cell of the extant clone.

Intestinal crypts contain multiple stem cells but are clonal populations because neutral drift recurrently eliminates all stem lineages except one (Lopez-Garcia, Allon M Klein, *et al*., 2010; Snippert *et al*., 2010). The clonality of human crypts has been previously inferred by a number of methods that use single or relatively few markers (Baker *et al*., 2014; Nicholson *et al*., 2018). The oscillator CpG sites represent a magnitude (>100 fold) increase in clock-like loci suitable for inferring recently-occurred stem cell dynamics. These oscillator CpG sites are common in methylation array data and show tissue specificity, likely reflecting differential gene expression between tissues (tick-tock sites are enriched at non-expressed loci). One of the major difficulties experiments with human tissue often encounter is the ‘snapshot’ nature of the data, making inference concerning dynamic processes difficult. To address this, we assessed how different temporal dynamics affect the *distribution* of methylation patterns across cells as measured in a ‘bulk’ sample consisting of many cells (such as an individual crypt) which, together with the relatively high *de novo* error rate of methylation, allowed us to track the stem cell dynamics within individual crypts. Oscillator CpG sites have different error rates and a key to analysis is to match error rates with the underlying rate of cellular dynamics. An oscillator that changes too fast fails to record cell dynamics because tick-tock methylation becomes desynchronized even in closely related cells. An oscillator that changes too slowly will maintain synchrony between distantly related cells and not capture more contemporary cell turnover. However, by matching CpG methylation oscillation rates with the biological interval of interest, we demonstrated the ability to infer the stem cell dynamics within glands. Stem cell numbers may have important fundamental roles in cancer risks because mutations that lead to cancer can only accumulate in long lived stem cell lineage (Ricci-Vitiani *et al*., 2007). Interestingly, consistent with experiments in mice (Kozar *et al*., 2013), we inferred only small differences in stem cell numbers between SI and colon crypts (SI crypts contains approximately 16% more stem cells than colon). Whereas colon carcinoma is the fourth most common human cancer (Siegel *et al*., 2021), SI carcinoma is between 14-50 times less common (Raghav and Overman, 2013; Siegel *et al*., 2021), even though their tissues have similar numbers of crypts and accumulate similar numbers of mutations during aging (Blokzijl *et al*., 2016). According to the “bad luck hypothesis” (Tomasetti, Li and Vogelstein, 2017), the discrepancy in cancer rates could be explained by differences in the stem cell dynamics of the two tissues, with more stem cells dividing more rapidly carrying a higher risk of progressing to cancer. We only detect moderate differences in the number of stem cells and the replacement rates per crypt between small intestine and colon. Hence, our data and analysis indicate that much lower SI carcinoma rates are unlikely to be solely attributable to the difference in stem cell dynamics between the two tissue types. We did observe a slight increase in the total number of replacements per crypt in non-dysplastic familial adenomatous polyposis (FAP) colon crypts that carry heterozygous *APC* mutations, perhaps suggesting that the “first-hit” loss of *APC* in the development of sporadic CRC confers a selective advantage, which may help explain why *APC* mutations are common in colorectal cancers.

We further demonstrate that CpG oscillators are present in hematopoietic cells and can be used to reconstruct clonal dynamics within the hematopoietic system. The identity of the oscillating CpG sites in hematopoietic cells differs from the epithelial oscillators, likely reflecting that oscillators tend to be found at non-expressed genes and the fact that gene expression patterns vary between tissues. Our blood studies illustrate the ability of oscillators to detect clonal hematopoiesis, with increases in average oscillator variances with clonality and characteristic W-shaped distributions in acute leukaemias. Chronic leukaemias had intermediate tick-tock variance increases and generally lacked W-shaped distributions, likely reflecting their slower growth and clinical persistence for years. Interestingly, there was a trend for an age-related increase in tick-tock variances that may reflect the increased incidence of clonal hematopoiesis of indeterminate significance or CHIP that becomes common with aging (Jaiswal and Ebert, 2019).

Our stem cell dynamics inference method relies on relatively inexpensive methylation arrays, but nevertheless a potential limit to this technique is the requirement of high-quality DNA derived from relatively small quantities of input material. The mathematical model-based inference necessarily relies on a number of assumptions (discussed in the methods) and the validity of these naturally affects the accuracy of our inference. Additionally, the dimensionality of the matrix encoding the stem cell dynamics scales quadratically with the number of stem cells, hence our inference framework is only tractable for reasonably small numbers of stem cells.

In summary, CpG methylation oscillator molecular clocks have many features ideal for the analysis of human cell populations. The oscillatory behavior of tick-tock CpG sites is otherwise elusive in polyclonal populations but becomes detectable in clonal cell populations. Oscillators can measure alterations that start or recur later in life and can infer changes that occur over a few years. Data are relatively easy to obtain with cheap, commercially available methylation arrays. Large numbers of potential tick-tock CpG sites suitable for the time intervals and cell populations of interest are found on these arrays. The ability to harness rapidly changing molecular clocks opens new opportunities to infer the histories of many different human somatic cell populations.

## Acknowledgments

AA and CG, TG and DS received support from the US National Institutes of Health National Cancer Institute (grant no. U54CA143970) and (U54 CA217376) respectively as well as supplemental support specifically for this collaboration through the Cancer Systems Biology Consortium (CSBC). AA, MT and RS also received funding from the US National Institutes of Health National Cancer Institute (U01CA232382) and from the Moffitt Center of Excellence for Evolutionary Therapy. TG received funding from Cancer Research UK (A19771 supporting EL). CG was funded by the BBSRC London Interdisciplinary Doctoral Programme (LIDo). RS is supported by the Wellcome Trust (grant no. 108861/7/15/7) and the Wellcome Centre for Human Genetics (grant no. 203141/7/16/7). SL funded by Wellcome Trust Senior Clinical Research Fellowship (206314/Z/17/Z) to SL. CB was funded by Wellcome Trust (209409/Z/17/Z). Core funding to the Wellcome Centre for Human Genetics was provided by the Wellcome Trust (090532/Z/09/Z). This research utilized the Cancer Research UK City of London High Performance Computing (HPC) facility, along with the Queen Mary’s Apocrita HPC facility, supported by QMUL Research-IT.

## Methods

### Methylation Array

Tissues were collected at the University of Southern California Keck School of Medicine from excess surgical samples taken in the course of routine clinical care, with Institutional Review Board approval. Crypts or endometrial glands were isolated using an EDTA-washout method as previously described (Yatabe, Tavaré and Shibata, 2001; Kim, Tavaré and Shibata, 2005). DNA methylation was measured with EPIC bead arrays (Illumina) using the using the Restore protocol and the manufacturers protocols (Illumina, 2011). IDAT files were processed with using the noob normalization function in the minfi R package (Aryee *et al*., 2014).

Blood methylation data were obtained from GEO (Edgar, Domrachev and Lash, 2002; Barrett *et al*., 2013) using beta values as provided. The data sets are GSE40279 (normal blood, Fig 6A (Hannum *et al*., 2013)), GSE73115 (ten-year serial samples, Fig 6B (Sierra, Fernández and Fraga, 2015)), GSE51759 (MDS (Zhao *et al*., 2014)), GSE42042 (ET, PRV, PMF (Pérez *et al*., 2013)), GSE106600 (CML (Maupetit-Mehouas *et al*., 2018)), GSE105420 (CMML (Palomo *et al*., 2018)), GSE62298 (AML (Ferreira *et al*., 2016)) and GSE69229 (ALL (Gabriel *et al*., 2015)).

RNA expression data for normal tissue derived from 40 patients was retrieved from The Cancer Genome Atlas (Muzny *et al*., 2012).

### Derivation of Oscillatory CpG loci

To isolate those CpG sites that behave in a clock-like fashion, it was first necessary to filter out those loci which are likely to have a regulatory function or change their methylation status over the length of the crypt. This was done by selecting only those CpG sites that lie in the ‘open sea’ (further than 4kb from a CpG island). Furthermore, probes of CpG loci that were identified (Chen *et al*., 2013; Pidsley *et al*., 2016) as being cross-reactive were filtered out, along with CpG loci positioned on sex-determinant chromosomes. Given the relatively low amounts of DNA contained within a single crypt, we also filtered out probes that were likely to have experienced incomplete binding by restricting our analysis to probes which had a total intensity greater than 1200 (arbitrary units).

The Illumina EPIC array features two different probe types, Type I and Type II (Pidsley *et al*., 2016). Type I probes feature a higher dynamic range, leading to the two probe types having different underlying distributions of beta values. Due to difficulties in simultaneously modelling the two different probe types, and given that Type I probes are overrepresented in CpG dense regions of the genome, the analysis was restricted to Type II probes.

### Mathematical Model of Methylation Within the Stem Cell Niche

We developed a stochastic model to describe how the fraction of methylated alleles (beta value) in the stem cell niche of a given CpG locus changes over time. This model draws on previous attempts (Lopez-Garcia, Allon M Klein, *et al*., 2010; Kozar *et al*., 2013) to model the behavior of the stem cell niche in colonic crypts, but with a number of modifications that account for the differences when considering methylation as a lineage tracing marker, rather than DNA. Namely, whilst DNA mutations occur relatively infrequently, allowing for a model that only considers a single mutant population expanding or contracting with reference to a single wild-type population, the relatively high methylation switching rate requires us to consider the potential of multiple clones existing simultaneously. Further, whilst DNA mutations can be generally regarded as irreversible, the methylation status of a given cell (that is, whether a particular cell is homozygously (de)methylated, or heterozygously methylated) can theoretically oscillate, necessitating a careful consideration regarding the possible ways the overall beta value can change.

For this reason, we made the simplifying assumption that the population was well-mixed, such that any of the *S* stem cells can replace any of the other *S −* 1 stem cells with equal probability, and that these replacements occur at a constant rate *λ* per stem cell. This assumption greatly simplified our analysis, as the system can be fully characterised using just 2 state variables: *k* – the number of stem cells containing a single methylated allele, and *m* – the number of stem cells containing 2 methylated alleles. The admitted states are constrained by the inequality 0 *≤ k* + *m ≤ S*, for a total of 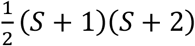 states.

Along with the replacement process, we assumed that a previously unmethylated CpG locus could spontaneously become methylated with a rate μ per year, and conversely, that a previously methylated CpG locus could spontaneously become demethylated with a rate γ per year.

To develop the series of ordinary differential equations that fully determine the system, we considered the ways in which a state (*k, m*) could transition to a state (*k*^′^, *m*^′^). As an example, if we consider Figure 1C, we observe that of the *S =* 5 stem cells, 3 of the stem cells are heterozygously methylated and 1 of the cells is homozygously methylated, hence the system is initially in state (*k =* 3, *m* = 1). To transition to state (*k*^′^ = 3, *m*^′^ = 2), the homozygously methylated stem cell must clonally expand, replacing the homozygously demethylated cell. The rate at which any one of the stem cells replaces another is *λS =* 5*λ*, but of the *S*(*S −* 1) = 20 possible transitions, only 1 would lead to the desired (3, 2) state, hence the rate at which the system transitions (3, 1) → (3, 2) is 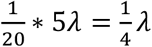. We continue this process (see the supplementary information), considering the general transition (*k, m*) → (*k*^′^, *m*^′^), deriving the master equation:

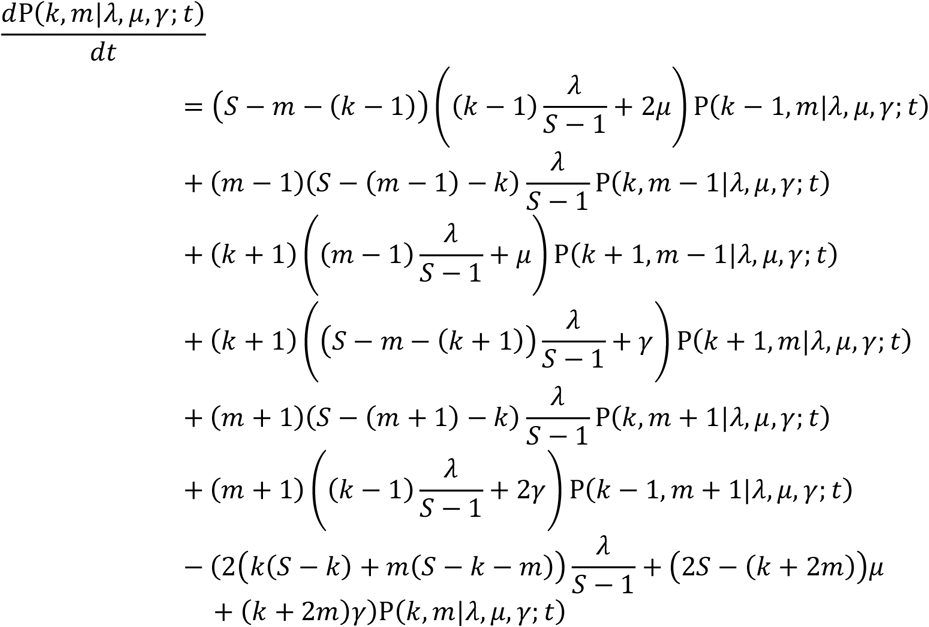

This linear series of differential equations can be solved computationally by rewriting the equations into a matrix equation, 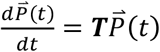, and applying matrix exponentiation to the resulting transition matrix, ***T***.

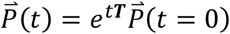

Given that all the stem cells within a niche are initially clonal, we assumed that it was equally likely to find a given CpG locus as homozygously methylated or unmethylated across all the stem cells within the niche at time 0.

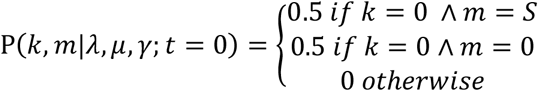

However, the methylation status of individual cells is not available using methylation arrays, hence the hidden states must be marginalized over to calculate the probability of there being *z* methylated copies within the stem cell niche (note that 0 *≤ z ≤* 2*S*). This can be achieved by summing the various combinations of *k* and *m* states that satisfy the equation *z = k* + 2*m*.

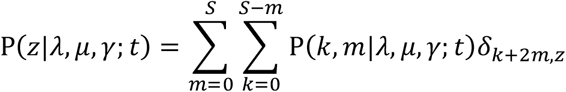

The resulting distribution of p(*z*|*λ, μ, γ*; *t*) can qualitatively reproduce the characteristic W-shape exhibited in the methylation fraction of individual crypts.

In developing our mathematical model describing the stem cell niche of intestinal crypts, a number of simplifying assumptions were made. Firstly, the assumption of a well-mixed population differs from Lopez-Garcia *et al*.’s (Lopez-Garcia, Allon M Klein, *et al*., 2010) assumption of a ring geometry; Lopez-Garcia *et al*.’s model considers the clonal expansion or retraction of a single clone, whereas our model must account for the possibility of multiple clones due to the increased mutation rate of the epigenome. As such, the assumption of a well-mixed population was chosen to minimize the mathematical complexity of the resulting model. Whilst it is true that LGR5+ cells likely reside in a roughly ring like structure, previous findings (Ritsma *et al*., 2014) that murine stem cells can exchange places within the niche lends support to treating the population as well mixed.

Furthermore, we assumed that the replacement rate, methylation rate and demethylation rate are constant over a patient’s lifetime. Whilst previous research indicates that the stem cell division rate lowers over a patient’s lifetime (Tomasetti *et al*., 2019), and our findings are consistent with such a decrease, it is likely that both the replacement rate and the methylation error rate are proportional to the cell division rate, such that the ratio of the two does not change over time. In this way, our model describes the stem cell dynamics of an individual crypt, averaged over a patient’s lifetime.

### Error Model

The probability distribution calculated above, p(*z*|*λ, μ, γ*; *t*), gives the probability that exactly *z* of the 2*S* alleles (across *S* stem cells) are methylated at a particular CpG locus; however, the Illumina EPIC array quantifies the methylation level at specific loci aggregated over the whole crypt. As such, we introduced an error model to link the measured *β*-value with the ‘true’ *z*-value at a specific site. We chose to model the measured *β* values as a mixture of *z* beta distributed random variables, each with a mean value determined by *z* and a scale parameter *k*_*z*_.

To account for the background noise of the array, the mean value of each beta peak was set to be equal to a linear transform of 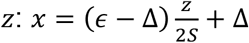, with the parameters describing this transform (*ϵ* and *Δ*) to be inferred. The scale parameters (sometimes referred to as the sample size),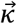, of each beta peak were modelled as hierarchical, with each *κ*_*z*_ being drawn from a lognormal distribution parameterized in terms of the population mean, *θ*, and its standard deviation, *σ*. These hyperparameters were also inferred during the Bayesian inference.

### Likelihood and prior functions

As rate parameters are naturally positive quantities, *λ, μ* and *γ* were constrained to positive real values by defining the prior distributions in terms of positive half-normals with a scale informed by prior literature. Following Nicholson *et al*.’s (Nicholson *et al*., 2018) finding that the replacement rate is approximately 1.3 replacements per stem cell per year, we set the scale of the prior on the replacement rate equal to 1. Similarly, *θ* and *σ* were also constrained to positive values using broad half-normal prior distributions, with a scale of 500 and 50 respectively. The lognormal hierarchical prior distribution naturally constrains 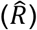 to real values. The “offsets” in the linear transform, *Δ* and *ϵ*, were constrained to lie between 0 and 1 by placing a beta distribution on each parameter, such that the mean prior value was and 0.95 respectively.

The behavior of individual CpG loci was assumed to be independent, such that the likelihood of all *N =* 1794 CpG loci was the simply the product of the per-CpG likelihood, computed according to the mathematical model outlined above.

Likelihood:

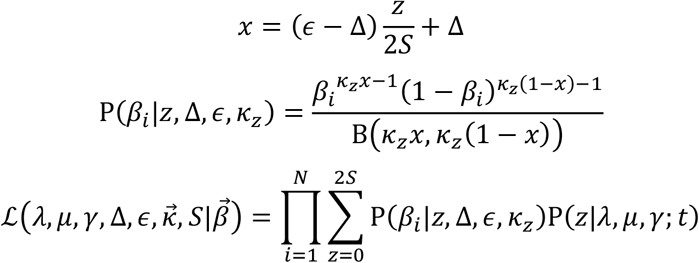

Hyperpriors:

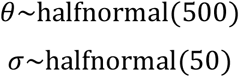

Priors:

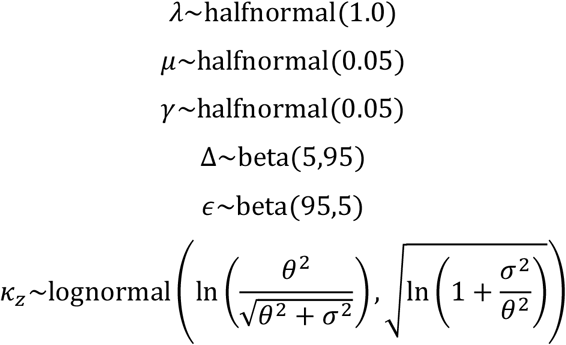

### Bayesian inference

A Bayesian inference methodology was developed to infer the biological model parameters (number of stem cells within the stem cell niche (*S*), replacement rate per stem cell per year (*λ*), and methylation (*μ*) and demethylation (*γ*) rate per CpG locus per stem cell per year) directly from the distribution of oscillatory beta values for each crypt.

Investigation of simulated datasets revealed that the resulting posterior distributions were multi-modal, with each *S* value associated with a local-maxima (due to the correlation in the posterior between *S* and *λ*). This multi-modality can make the posterior difficult to explore using traditional MCMC techniques, such as Hamiltonian Monte Carlo. To overcome this, a nested sampling method (Skilling, 2006) was developed to calculate the Bayesian evidence (marginal probability density, *Z*) of each *S* value considered (*S* ∈ [3. .20]) whilst simultaneously generating samples from the posterior associated with each value of *S*. The probability of *S* for a given crypt can then be calculated as:

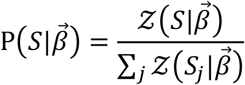

The full posterior can be approximated by drawing samples from each *S* mode with a weight equal to the inferred probability of *S*. The nested sampling was performed using dynesty (Speagle, 2019), a python implementation of the nested sampling algorithm, using the ‘rwalk’ sampling option, such that new live points are generated from existing live points under random walk behavior.

To ensure that the posterior samples had converged to the equilibrium distribution, four independent samples were run with random initializations for each sample, and the rank-normalized potential scale reduction statistic (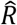) calculated (Gelman and Rubin, 1992; Vehtari *et al*., 2020). 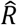 was found to be less than 1.1 (a typical threshold used to determine convergence) in all cases. The inference code can be obtained from https://github.com/CalumGabbutt/ticktockclock.git

### Tissue-specific differences in stem cell dynamics

To compare the stem cell dynamics of different tissue and disease types in a statistically rigorous manner, we must account for the hierarchical patient structure (that is, we have multiple glands from each patient which are likely to be correlated) whilst controlling for the age and sex of each patient. We developed a hierarchical Bayesian generalized linear model (GLM) using a log-link function to constrain our dependent variable to be positive (presented fully in the supplementary material), and take a hypothesis testing by parameter estimation approach (that is, the difference between small intestine and colon is statistically significant if the 95% equal-tailed credible interval excludes 0).

### Spatial Model of the Crypt

A crypt ignoring villi in the small intestine, forms a cylindrical geometry with stem cells at the base and a crypt wall moving up the crypt. Here we have developed an off-lattice mechanistic agent-based model of the human crypt using the HAL modeling framework (Bravo *et al*., 2020) capable of representing a crypt of the small intestine or colon (Fig. 3D). The cylindrical unit is separated into two compartments, the stem cell compartment represented as a pool at the base of the crypt and then the wall of the crypt where transient amplifying cells are pushed upwards until they are removed from the top of the crypt. The spatial model of the crypt is dynamic in the sense that the *x* and *y* dimensions are calculated using the total populations size (*N*_*T*_) and the stem cell pool radius (*Ψ*). The *x* dimension is defined as *x =* 2*πΨ*. The center of the stem cell pool is placed such that the origin of the center of this circular stem cell pool whose size, and thus number of stem cells allowed within this pool, is placed at (*h, k*) where *h = x/*2 and *k = Ψ* + 5. Division for each stem cell is defined by *ρ*_*c*_ which is randomly assigned as the hourly cell cycle defined by *p*_*c*_*∼U*(*ρ*_*min*_, *p*_*max*_) where *ρ*_*min*_ and *ρ*_*max*_ are *ρ* ± 4 hours.

As a cell approaches *ρ*_*c*_ the cells diameter doubles for the five hours/timesteps preceding the cells division. Upon division, both daughter cells occupy this space. When a stem cell, defined by *d*(*x*_*c*_, *y*_*c*_) *≤ Ψ* where 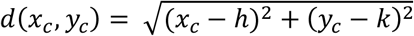, divides the daughter cells can be placed in any arrangement around the parent cells *x*_*c*_ and *y*_*c*_ position, differentiated cells can only be placed vertically (i.e. the *x*_*c*_ values are equal). The base of the crypt wall is set just above the origin of the stem cell pool plus *Ψ* and a small offset to provide space so that no cell forces interact between the stem cell pool and the base of a stem cell wall. If *d*(*x*_*c*_, *y*_*c*_) *> Ψ* then the cell is moved to the base of the stem cell wall where the cells new position (*x*_2_, *y*_2_) is given as *y*_2_ and *x*_2_ is given by the cells exit radians, *rad*_*s*_, given by *atan*2(*y, x*) so that the cells position along the *x* dimension is 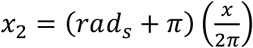 Boundary conditions for the cells within the crypt wall are periodic (i.e. allowed to wrap around) and no-flux at the top and bottom of the crypt (i.e. no cell can breach these boundaries). A run step in the model is hourly and updates to cell positions occur for the whole crypt are applied each timestep. We give each cell 1794 CpG loci (with the possible status of 0 for de-methylated or 1 for methylated). At each division these loci can switch methylation status at a rate defined by *ω* upon division.

At each hourly time step we assume that the forces acting on each individual cell to be at equilibrium, 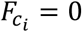, where 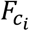is equal to the contact force between cell *i* and its neighbors. For two cells whose radius is *R*_*i*_ and *R*_*j*_, respectively, their contact force between them is based on a linear spring contant model (Hooke’s law) and is calculated as

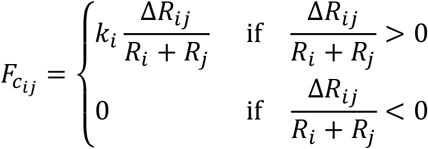

Assuming that each cell has the same spring constant *k*, the overlap of cells 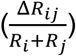, and the overall number of cells in contact with any given cell (*n*_*i*_) gives the velocity for an individual cell is 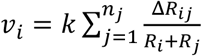. The modelling framework can be obtained from https://github.com/MathOnco/ticktockspatialmodel.git.

### Inference of Stem Cell Numbers on the Spatial Model

In order to provide insights into the oscillatory signal from a first principles model of the homeostatic crypt (balanced birth/death with a methylation error rate) we have to add noise to the output data of the spatial model. This is because the inference framework is designed to fit the noisy experimental data and that oscillatory CpG sites with values of zero or one are not captured within the data. In order to add a small amount of noise to the output of the perfect beta distributions output by the spatial model a binomial is used with two offsets to provide a distribution that the inferences can be performed on. For each beta value a sample size (*κ*) of 1000 is taken from a beta distribution using an offset from zero (*Δ =* 0.04) and an offset from one (*η* = 0.92) (Fig. 3D). The script required to add noise to this model is accompanied with the inference framework (see add_noise.py). Once the beta values with noise are added the inference framework is executed for each model simulation’s beta value distributions for across stem cell number ranges from 2-9, 3-10 and 8-15 respectively using 400 live points for the dynesty sampler (Speagle, 2019).

## Supplementary Information

### Supplementary Figures

**Supplementary Figure 1:**
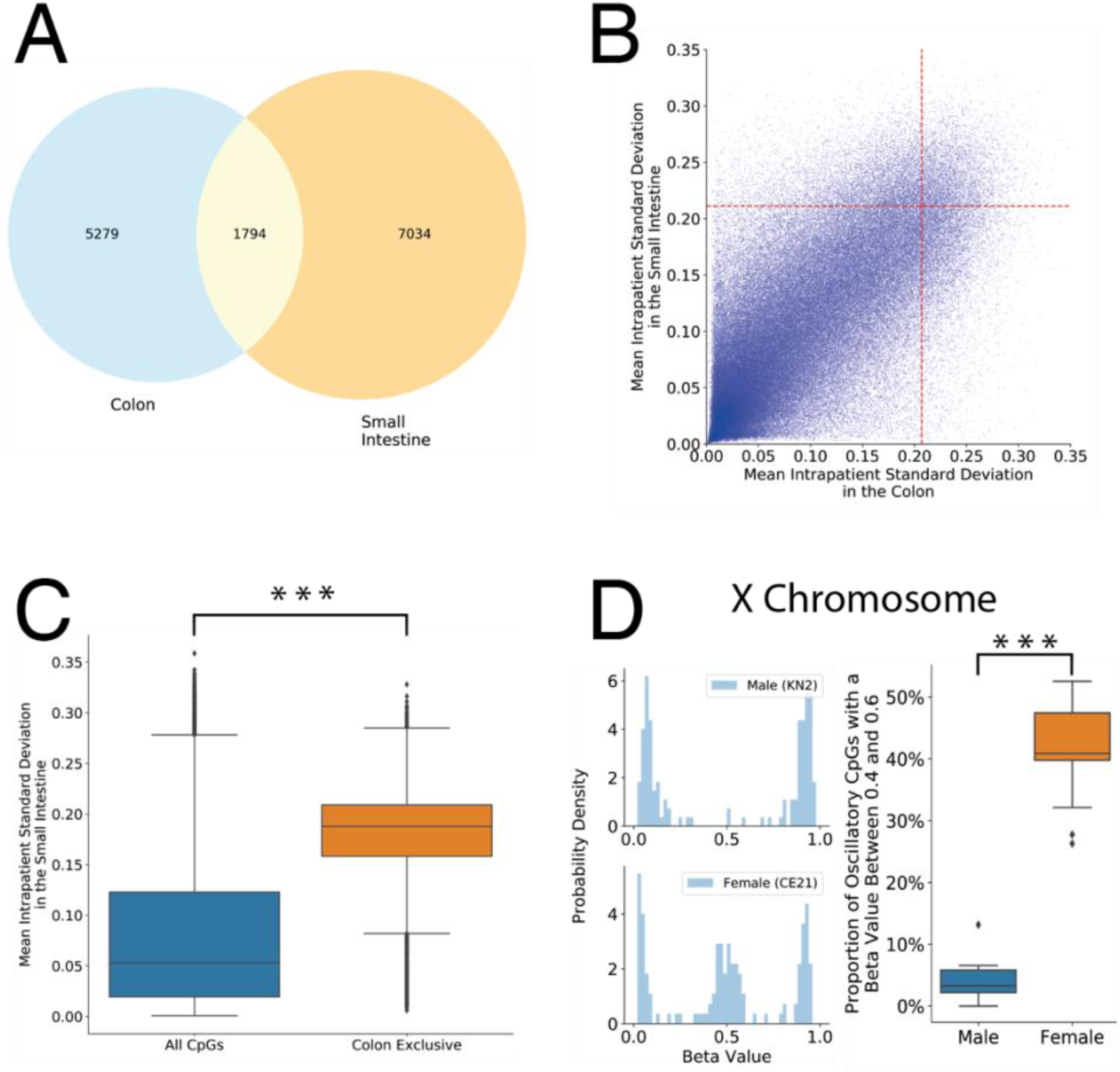
Additional Analysis of Oscillatory CpG Identification Process. **A:** Venn diagram showing the overlap of CpG loci identified as oscillatory in the colon and the small intestine. **B:** Scatter density plot (with the density plotted on the log-scale) of the heterogeneity metric (mean intra-patient standard deviation) of CpGs in the colon and small intestine, with the cutoff of the top 5% most heterogenous loci indicated in red. **C:** Comparison of the heterogeneity metric of the colon exclusive oscillatory CpG loci (i.e. those identified in the colon but not the small intestine) to all type II CpGs, within the small intestine samples. The colon exclusive CpG loci are significantly more variable (p<0.001, Mann Whitney U test). **D:** An extension of the oscillatory CpG identification process to CpG loci located on the X chromosome. We present example tick-tock distributions for these X-chromosome CpG loci for a male and a female crypt, confirming the predictions from theory that the male crypts lack the peak at 50% as they contain only a single copy. To test whether this relationship holds in general, we compare the proportion of tick-tock CpG’s with an intermediate beta value (0.4 *≤ β ≤* 0.6) between all colon crypts from males and females, confirming that males have a significantly lower probability mass near 50% (p<0.001, Mann Whitney U test).

### Derivation of Model Describing Methylation Within the Stem Cell Niche

Consider a single CpG locus within a fixed population of *S* stem cells. Within each stem cell, the locus is assumed to be diploid, so each stem cell contains 2 alleles at this locus. In this way, there are 3 possible “states” for a given stem cell, (i) neither allele methylated, (ii) both alleles methylated, (iii) or one allele methylated whilst the other is unmethylated. We are interested in the population methylation level, so we assume that the population is well-mixed, which allows us to characterize the system using just 2 state variables: *k* – the number of stem cells containing a single methylated allele, and *m* – the number of stem cells containing 2 methylated alleles. The number of stem cells containing 0 methylated alleles is then given by *S − m − k*.

The states are constrained such that 0 *≤ k, m ≤ S* and *k* + *m ≤ S*, which allows us to calculate the total number of possible states by considering all possible combinations of *k* and *m*. If we first consider the case when *m =* 0, then *k* can take any value between 0 and *S* giving a total of *S* + 1 states. If we next consider the *m =* 1 case, then *k* can take any value between 0 and *S −* 1, a total of *S* states. We can continue in this fashion for each of the *S* + 1 possible states for *m*, such that the total number of states is

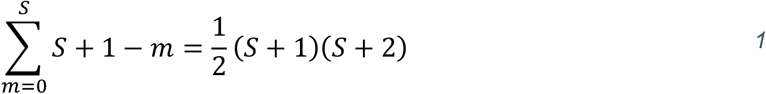

We assume that there are three possible processes that can change the population methylation level (*k, m*) → (*k*^′^, *m*^′^):

1. an unmethylated allele spontaneously becoming methylated (which, for a single unmethylated CpG locus, occurs at a rate μ per allele per stem cell per year)
2. a methylated allele spontaneously becoming unmethylated (which occurs at a rate γ per allele per stem cell per year)
3. one stem cell replacing one of the other *S −* 1 stem cells (which occurs at a rate *λ* per stem cell per year).

To formulate a system of differential equations that characterize the rates at which the population methylation changes, we first consider the probability the system in state (*k, m*) at time *t* transitions to state (*k*^′^, *m*^′^) within the time *t* + *δt* (where we assume *δt* is small enough that the probability of a “double-jump” is negligible).

If we are in state (*k, m*), then the probability that one of the *k* heterozygous methylated stem cells becomes unmethylated (via process (2)) in a time period *δt* is:

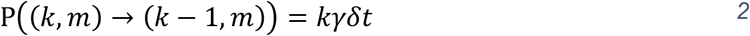

And the probability that the one of the *m* homozygous methylated stem cells (representing 2m methylated alleles) undergoes process (2) is:

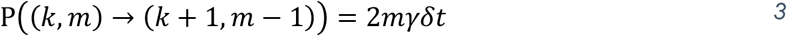

Similarly, considering methylation (process (1)), there are a total of 2*S − k −* 2*m* unmethylated alleles where the process could occur. The probability that one of the homozygous S-k-m unmethylated stem cells becomes heterozygous is:

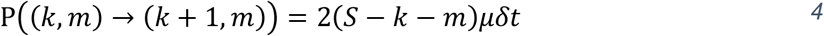

And the probability that one of the heterozygous methylated stem cells becomes homozygous methylated is:

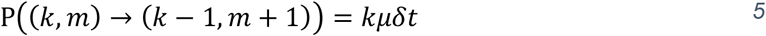

Let us now consider the replacement process. In a time period *δt* the probability that a replacement occurs is *Sλδt*. There are *S*(*S −* 1) possible replacements: *S* possible cells that can expand, which must replace any of the *S −* 1 other cells. To go from state (*k, m*) to a different state (*k*^′^, *m*^′^), we require the expanding stem cell to replace a cell with a different methylation status. Therefore, the probability of the transition (*k, m*) → (*k*^′^, *m*^′^) is equal to the probability that any of the cells replaces another, *Sλδt*, multiplied by the number of ways that particular transition could occur, and normalised by the total possible number of transitions.

To give a concrete example, consider the stem the cell niche illustrated in Figure 1C, which contains 5 stem cells and is initially in the state (*k =* 3, *m =* 1). There are a total of 5 *\** 4 = 20 possible replacements. Clearly, if one of the heterozygous stem cells replaces another of the heterozygous stem cells, the population methylation level will not change. To jump to the state (*k =* 3, *m =* 2) as illustrated in Figure 1C, only one replacement (the homozygous methylated stem cell replacing the homozygous unmethylated stem cell) allows the specified jump, hence the probability of the jump (3,1) → (3,2) in the time *δt* is 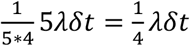. To generalise this, the fraction of possible transitions that give rise to the particular jump (*k, m*) → (*k*^′^, *m*^′^) is equal to the multiplicity of the expanding cell multiplied by the multiplicity of the replaced cell, divided by *S*(*S −* 1).

Applying the same logic, we can derive the probability of all six possible state transitions via replacement:

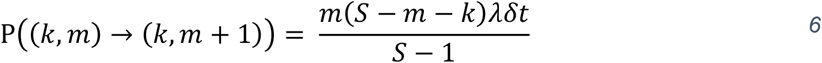

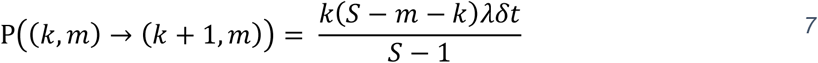

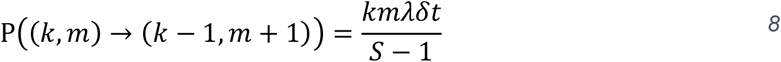

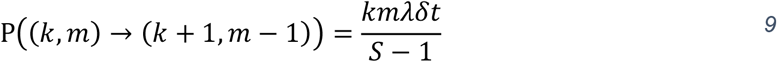

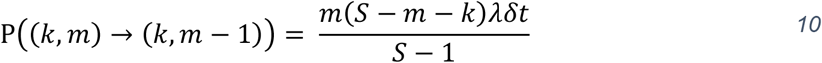

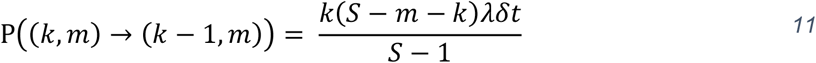

The methylation switching and replacement processes that we have considered separately above are independent, allowing us to simply add the probabilities together (again, assuming that *δt* is small enough that the probability of two processes occurring in *δt* is negligible) to find the total probability that a given transition would occur:

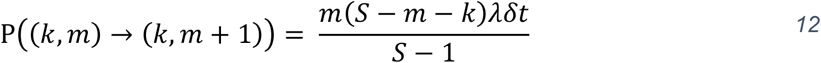

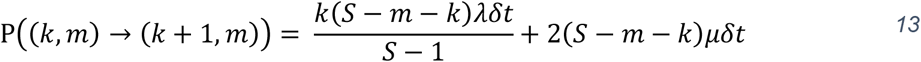

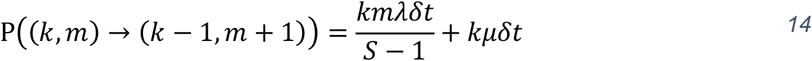

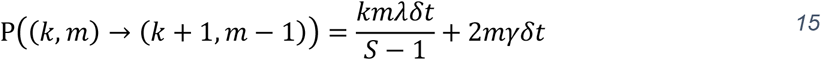

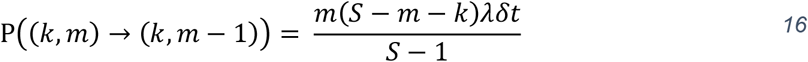

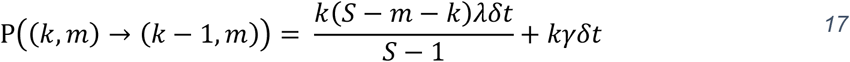

We have considered above the transitions leading “out” of the state (*k, m*) into adjacent states (*k*^′^, *m*^′^). However, we can also consider the jumps “into” the state (*k, m*) from the adjacent states (*k*^′^, *m*^′^):

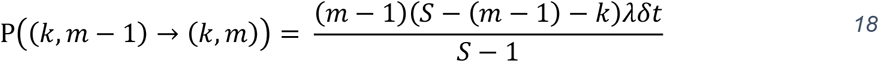

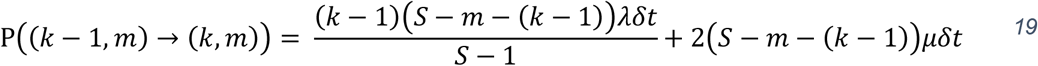

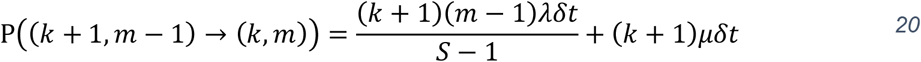

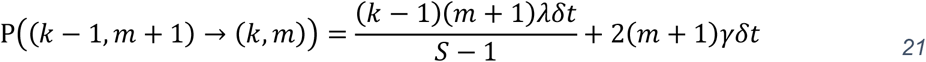

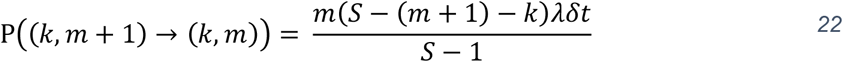

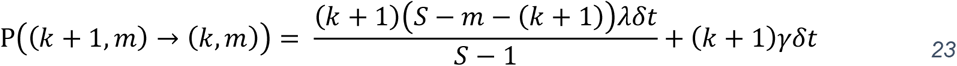

So far, we have considered the probability that the system changes from state (*k, m*) to state (*k*^′^, *m*^′^) within time *δt*. However, we primarily want to know the probability of the system being in state (*k, m*) at time *t*, p(*k, m; t*), and how this changes over time. For the system to be in state (*k, m*) at time *t* + *δt*, either (i) the system must have been in state (*k, m*) at time *t* and has not transitioned out of the state (which is equal to 1 minus the probability of transitioning to an adjacent state, defined by equations 12-17), (ii) or the system was in a different (adjacent) state (*k*^′^, *m*^′^) at time *t* and has transitioned into the state (*k, m*) in time *δt* (defined by equations 18-23):

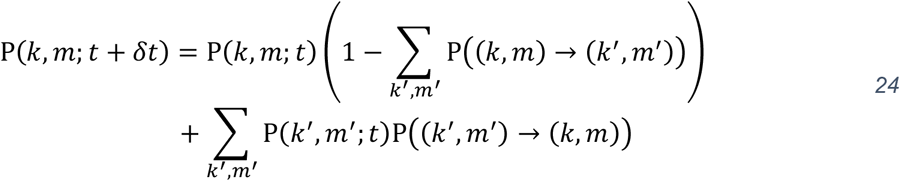

We can rearrange equation 24, factoring out the common factor of *δt* in the p((*k*^′^, *m*^′^) → (*k, m*)) terms and take the limit *δt* → 0:

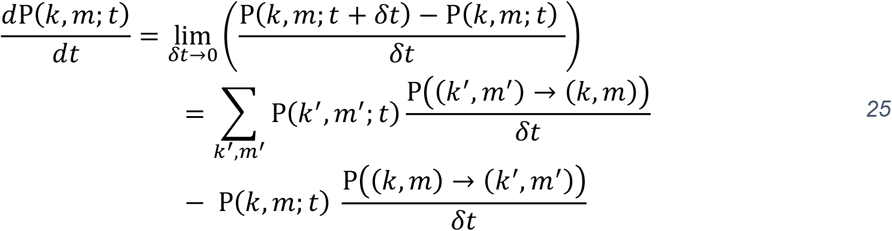

The sum over equation 12-17 in the final term evaluates as:

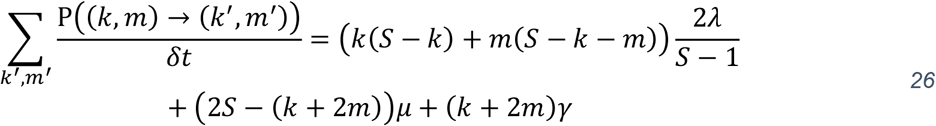

Due to the constraints on *k* and *m*, we consider the differential equations for (*k =* 0, *m =* 0), (*k = S, m =* 0) and (*k =* 0, *m = S*) separately. Combining equations 25, 26 and 18-23, we derive the following set of differential equations:

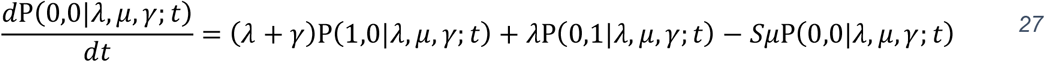

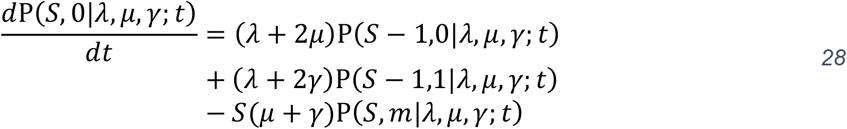

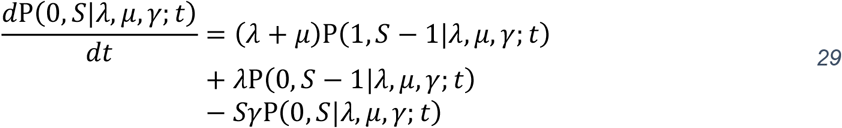

Otherwise:

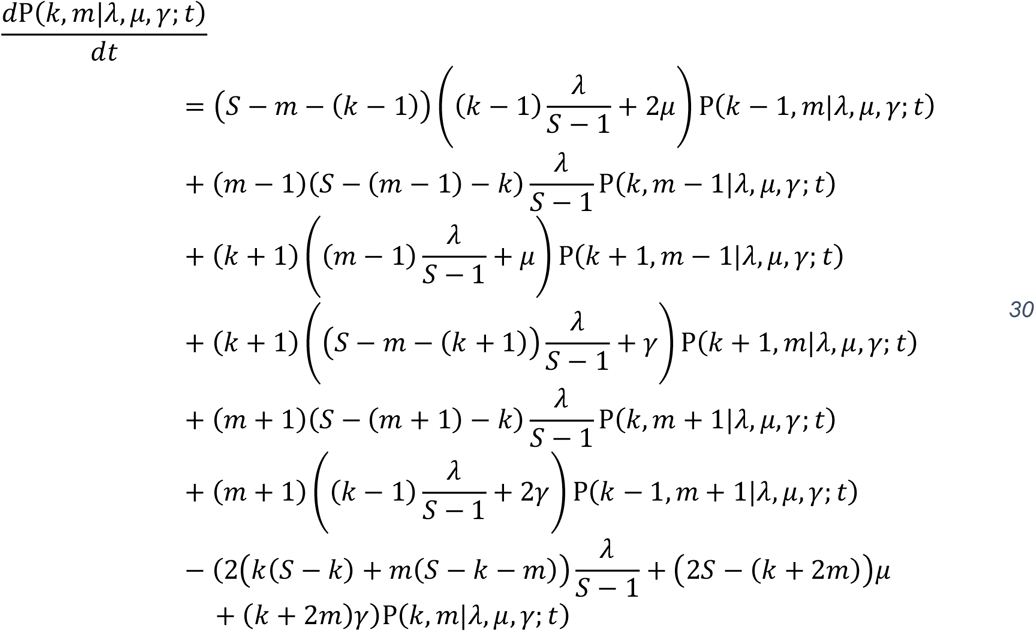

This master equation determines how the methylation statues of a single CpG locus within the stem cell niche evolves over time. The replacement, methylation and demethylation rate are assumed to be constant, hence this process is Markovian and we are able to solve this using standard matrix exponentiation.

### Bayesian Analysis of the Effect of Tissue Location and Disease State on Stem Cell Dynamics

The Bayesian pipeline described in the main body of the text allowed the posterior distribution of the parameters defining the stem cell dynamics (namely, the effective number of stem cells, *S*, and the replacement rate per stem cell, *λ*) of each individual crypt to be inferred. To interrogate the effect of age, sex, tissue location (colon, small intestine and endometrium) and the disease state of colonic crypts (AFAP/FAP) on stem cell dynamics, we take the posterior mean of *S* and *λ* as representative of the inferred distribution for each crypt.

As a matter of notation, let there be *K* patients subscripted with *k = [*1.. *K]* and *N* crypts subscripted with *i = [*1.. *N]*. The age of the *k*^*th*^ patient is *t*_*k*_, which we normalise to be between 0 and 1 by dividing each patient’s age by the maximum age in the patient cohort. Similarly, the sex the *k*^*th*^ patient is encoded as a dummy variable, which equals 0 for female patients and 1 for male patients. The location/disease state of each crypt is encoded with the dummy variables *x*_*i,j*_ for *j ∈ {Colon, Small Intestine, FAP, AFAP, Endometrium}*.

We fit the parameters determining stem cell dynamics *y = {S, λ}* using a generalised linear model with a gamma-distributed dependent variable (this accounts for the fact *S* and *λ* are strictly positive). Let *y*_*i,k*_ be the dependent variable with expectation ŷ_*i,k*_, then we employ the natural log as a link function *g*(*ŷ*_*i,k*_) *= ln*(*ŷ*_*i,k*_). *y*_*i,k*_ is then gamma distributed with mean ŷ_*i,k*_ and a tissue/disease-specific standard deviation *ϕ*_*j*_.

We use the parameterization of the gamma distribution in terms of its shape (*Ψ*) and rate (*ω*):

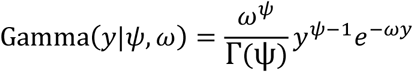

The mean of this distribution is 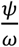 and the variance is 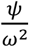. Hence, to parameterise the gamma distribution in terms of its mean (ŷ) and standard deviation (*ϕ*), we apply the transformation 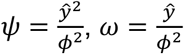

Our dataset contains multiple samples from the same patient, so we assume the offset in the linear predictor is drawn for each patient from a hierarchical normal distribution with mean *ν* and variation *σ* (hence accounting for random inter-patient variability, not attributable to the factors we are explicitly modelling). Similarly, to maximize the information that can be drawn from the data, we allowed the tissue/disease-specific intrapatient standard deviation, *ϕ*_*j*_, to be drawn from a lognormal distribution, with a population mean *ρ* and standard deviation *ζ*.

Priors:

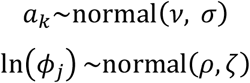

Model:

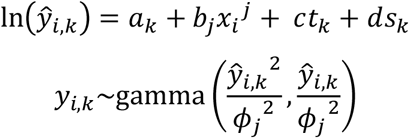

The hierarchical Bayesian model was fit to the data using pystan, a python implementation of Stan (Carpenter *et al*., 2017), a probabilistic programming language that allows for rapid MCMC sampling.

Because a log-link function was used to ensure the positivity of *ŷ*_*i,k*_, encode the difference between each tissue-type or disease-state, and colon on the log scale. We take the exponential transform of each of these regression coefficients to derive the posterior for the relative stem cell number and replacement rate of each tissue-type and disease-state relative to colon. We take a hypothesis testing by parameter inference approach, where the effect of a particular tissue/disease on the dependent variable is termed significant when the 95% equal-tailed credible interval does not overlap 0. The hierarchical Bayesian model that we have developed naturally penalizes increasing numbers of parameters, hence there is no need to apply a multiple test (Gelman, Hill and Yajima, 2012).

### Whole Blood Simulations

Whole blood was simulated in Java using the HAL framework (Bravo *et al*., 2020) as a non-spatial agent-based model using 27,634 oscillating CpG sites as measured in the experimental data. Parameters (Table 1) for normal hematopoiesis are numbers of hematopoietic stem cells (N, HSCs), number of possible division events (T), CpG error rates (S, methylation and demethylation) for the oscillators, and HSC replacement dynamics (R). To model clonal expansion, a single cell is selected to grow upon induction, and added parameters are its expansion rate (E) and its final blood frequency of the clonal expansion (M). These clonal expansions result in the overall population size to grow until the appropriate final blood frequency is reached. The output of the simulations provides the beta values from the oscillating CpG sites and the overall distribution’s variance over time.

The number of HSCs was set at a lower value of 1000 initiating cells. This is much lower than the 30,000 based on the large number of HSC inferred by DNA sequencing studies (Lee-Six et al, 2018; Watson et al., 2020); however, the results shown here are invariant to more than 100 initiating cells. CpG error rates varied between CpG sites and were assigned based on the distribution averages of the 656 normal individuals from GSE40279. We found that some of the whole blood oscillators did not appear to have equal methylation and demethylation error rate because their averages tended to always be above or below 50% in multiple individuals. Hence, to better model and match the data, we used a look-up distribution table in the simulations in order to initialize cell’s oscillatory CpG parameters, with lower and unequal error rates at CpG sites with average methylation typically found near 0.4 (demethylation > methylation) or near 0.6 (methylation > demethylation) to maintain the variance of the 27,634 oscillators around 0.1 during cell divisions. The error rates varied between 0.0001 to 0.001 changes per division, with the highest error rates and more equal methylation and demethylation rates at CpG sites near 50% methylation.

Cell survival was set at exact replacement (one cell produces one living offspring), and results did not vary much if random replacement was simulated. A proportion of cells, no matter if it’s a founding disease cell undergo replacement at each timestep (Supplemental Table 1). For the neoplastic simulations in Fig 6D in the manuscript, the expansion rate (E) was varied to model either rapid expansion (visible or more than 5% leukemic cells within 1 year or 200 divisions) akin to acute leukemia, modest expansion (visible within 4 years or 12,000 divisions), or very slow expansion (visible within 6 years or 18,000 divisions). The extent of blood involvement was varied between 20% (black lines), 50% (blue lines) and 90% (red lines). These simulations indicate that how clonal expansions change whole blood oscillator variances depends both on how fast the expansion grows and to what extent it involves the blood. Rapid growth to high levels like acute leukemias results in high oscillator variances and characteristic W-shaped tick tock distributions. Slower growth to lower levels like chronic leukemias results in low oscillator variances and broader distributions that lack the W-shape. Interestingly, very indolent clonal expansions which may occur with CHIP (Jaiswal and Ebert, 2019) can result in small increases in oscillator variances, which may account for the age-related increase in oscillator variances seen in Fig 6A in the manuscript.

More sophisticated modelling with a better selection of whole blood oscillators could improve the extraction of ancestral information. For example, a selection of slower oscillators may improve the detection and analysis of indolent clonal expansions, where many of the faster oscillators return to average ∼50% methylation by the time the expansion reaches detectable blood levels.

The simulation framework can be obtained, along with sample simulation results, on GitHub through https://github.com/MathOnco/ticktockblood.git. A GUI compatible with most operating systems is accompanied to allow for rapid evaluation of different parameters.

Parameters used in simulations describing how the methylation distribution of well-mixed hematopoietic stem cells (HSCs) changes in response to the expansion of a single clonal population.

**Supplementary Table 1:** Parameters of Whole Blood Simulations

| Parameter | Description | Values |
| --- | --- | --- |
| $N$ | Number of HSCs, initial population | 100 |
| $T$ | Simulation time | 2,500 |
| $S$ | CpG error rates | 0.0001-0.001 per division |
| $\lambda$ | Cell survival, exact replacement | 0.6 |
| $E$ | Disease expansion rate | (0.1, 0.005, 0.00225) |
| $\omega$ | Final blood frequency | (0.2, 0.5, 0.9) |

